# eIF3b and eIF3i relocate together to the ribosomal subunit interface during translation initiation and modulate start codon selection

**DOI:** 10.1101/453688

**Authors:** Jose L. Llácer, Tanweer Hussain, Jinsheng Dong, Yuliya Gordiyenko, Alan G. Hinnebusch

## Abstract

During eukaryotic translational initiation, the 48S ribosomal pre-initiation complex (PIC) scans the 5’ untranslated region of mRNA until it encounters a start codon. We present a single particle electron cryomicroscopy (cryo-EM) reconstruction of a yeast 48S PIC in an open scanning-competent state in which eIF3b is observed bound on the 40S subunit interface. eIF3b is re-located with eIF3i from their solvent-interface locations observed in other PIC structures; however, eIF3i is not in contact with the 40S. Re-processing of micrographs of our previous 48S PIC in a closed state using currently available tools reveal a similar re-location of eIF3b and eIF3i from the solvent to subunit interface. Genetic analysis indicates that high fidelity initiation in vivo depends strongly on eIF3b interactions at the subunit interface that either promote the closed conformation of the PIC on start codon selection or facilitate subsequent relocation back to the solvent side of the 40S subunit.

## Introduction

Eukaryotic translation initiation is a complicated process that involves a complex of the 40S subunit and initiation factors binding to the 5′ end of mRNA and scanning the mRNA until a start codon is encountered in the ribosomal P site (Hinnebusch, 2017). First, the factors eIF1, eIF1A and eIF3 bind to the 40S, which facilitates the recruitment of methionyl initiator tRNA (Met-tRNA_i_) as a ternary complex (TC) with eIF2-GTP. eIF5, a GTPase activating protein (GAP) for eIF2, may be recruited along with TC or in complex with eIF3. The 43S preinitiation complex (PIC) thus formed is recruited to the capped 5′ end of mRNA by the eIF4 group of factors (Pelletier and Sonenberg, 2019) (Mishra et al., 2019). This 48S PIC, in an open conformation with the tRNA_i_ not fully engaged with the P site (P_OUT_ state), then scans the mRNA until a start codon is encountered. GTP bound to eIF2 can be hydrolysed during the scanning process, however, the phosphate (P_i_) product remains bound. Recognition of the start codon leads to a conformational change in the PIC to form a scanning-arrested closed complex, with Met-tRNAi more tightly bound (P_IN_ state), accompanied by the release of eIF1 and attendant dissociation of P_i_ (Hinnebusch, 2014; Hinnebusch, 2017; Aylett and Ban, 2017).

Among the eIFs, eIF3 is the largest and contains the greatest number of subunits, and is involved in every step of translation initiation, including TC and mRNA recruitment to the PIC, impairing the association of the 40S and 60S subunits, and modulating the fidelity of start codon selection (reviewed in Valasek, 2017, Nucleic Acid Research, 45, 10948-68). In mammals, eIF3 contains 13 subunits (a-j); however, in *Saccharomyces cerevisiae,* it is a much smaller with only 6 subunits (including the non-essential j/HCR1 subunit). The five essential subunits in *S. cerevisiae*, namely, a/Tif32, b/Prt1, c/Nip1, g/Tif35 and i/Tif34, comprise a conserved core complex found in all organisms, that interacts with the 7 other subunits found in mammalian eIF3. A recent study using advanced mass spectrometry revealed that the yeast eIF3 complex in solution adopts a globular 3D structure, with the WD40 domains/β-propellers of eIF3b and eIF3i, the RRMs of eIF3b and eIF3g and the N- and C-terminal ends of eIF3a and eIF3c all exposed to the surface. Interestingly, the PCI domains of eIF3a and eIF3c do not interact with each other in the free eIF3 complex (Zeman et al., 2019). This stands in contrast to the interaction between the two PCI domains observed in the structures of eIF3 present in various conformational states of the PIC, wherein the eIF3 complex seems to have undergone a large conformational change on binding to the 40S subunit (Hashem et al., 2013) (Aylett et al., 2015) (Llácer et al., 2015) (Llácer et al., 2018) (des Georges et al., 2015) (Eliseev et al., 2018). Indeed, eIF3 adopts an expanded conformation when bound to the PIC, in which the eIF3a/eIF3c PCI heterodimer occupies a position near the mRNA exit channel whereas the distinct module formed by the eIF3b, eIF3g, and eIF3i subunits (eIF3b-3i-3g) is positioned near the mRNA entry channel (Hashem et al., 2013) (Aylett et al., 2015) (Llácer et al., 2018) (des Georges et al., 2015) (Eliseev et al., 2018). The two modules are connected by the C-terminal (Cter) end of eIF3a.

A comparison of the structures of yeast and mammalian PICs harbouring eIF3 reveal that the five core subunits (a, b, c, g and i) occupy the same positions on the 40S subunit irrespective of the subunit complexity of eIF3. However, in the cryo-EM structures of partial yeast 48S PIC complexes in an open, scanning-competent state (P_OUT_; hereafter referred to as “py48S-open”) and in closed scanning-arrested state (P_IN_; hereafter referred to as “py48S-closed”) (Llácer et al., 2015), we observed a portion of the eIF3b-3g-3i module at an alternative location, positioned on the subunit interface near h44, uS12 and TC, where it could contribute to the 40S-60S anti-association activity of eIF3. Thus, a striking conformational change in eIF3 was observed in these py48S-open and py48S-closed complexes, in which eIF3 appears to encircle the entire 40S, compared to its configuration in the previously reported PICs where it is confined to the solvent-exposed surface of the 40S near the mRNA entry and exit channels.

In the late-stage of initiation, as observed in in the py48S-5N complex from yeast, the N-terminal domain of eIF5 (eIF5-NTD) replaces eIF1 on the platform, thereby replacing eIF1 interactions with tRNA_i_ that impede the P_OUT_ to P_IN_ transition with favorable eIF5-NTD/tRNA_i_ interactions that stabilize the P_IN_ state. Importantly, the eIF3b-3i-3g module is found on the solvent side of the 40S (Llácer et al., 2018) in the location observed previously in 43S PICs. This led us to hypothesize that the eIF3b/3i/3g module interacts initially with the solvent surface of the 43S PIC, relocates to the subunit interface at the onset of mRNA attachment or scanning up to the point of start codon selection, and then relocates back to the solvent side for the final steps of initiation. This second relocation step would be required for replacement of eIF1 at the P site with the somewhat bulkier eIF5-NTD, for dissociation of eIF2 from tRNA_i_ (owing to loss of eIF3b interactions at the subunit interface with eIF2γ), and finally for 40S-60S subunit joining (Llácer et al., 2018).

The positioning of portions of the eIF3b-3g-3i module at the 40S subunit interface in py48S-open and py48S-closed complexes showed how eIF3 can contact TC and eIF1 in the decoding center while remaining anchored through its other subunits/domains to the solvent side of the 40S. However, the portions of eIF3 bound on the 40S subunit interface were observed at lower local resolution compared to the rest of the PIC. In particular, our previously reported cryo-EM structure of “py48S-open”, at an overall resolution of 6.0 Å (Llácer et al., 2015), revealed well-resolved densities for eIF1, eIF1A, all three subunits of TC including eIF2β, and portions of eIF3; however, the PCI domains of eIF3a and eIF3c on the solvent surface were not observed. Hence, to reveal in greater detail the interactions of all five eIF3 subunits with the PIC, particularly those occurring at the subunit interface, we sought to obtain a map of the 48S PIC in an open conformation at higher resolution. This has been achieved here by collecting a larger data set with an improved sample (described below) and by employing image processing procedures that were unavailable earlier, including signal subtraction and 3D classification using masks (Bai et al., 2015), to obtain superior and more homogenous 3D classes.

Here we present a structure of the yeast 48S PIC in an open scanning-competent state with density for eIF3 PCI domains bound on the solvent side of the 40S and density for the quaternary complex of eIF3b-3i-3g-3a-Cter bound on the subunit interface of the 40S. The structure reveals that the entire eIF3b-3i-3g-3a-Cter module has been relocated from its position on the solvent side, observed in other yeast and mammalian PIC structures, to the subunit interface by virtue of an extended conformation of eIF3a-Cter. This finding is in contrast to our earlier report on the py48S-open complex where only a portion of the eIF3b-3i-3g-3a-Cter module appeared to be present at the subunit interface. In our new structure, the eIF3b β-propeller binds near h44 on the 40S subunit interface and interacts with TC, and the eIF3b RRM contacts eIF1 bound near the P site, while the eIF3i β-propeller makes no direct contact with the 40S. A similar position of the eIF3b-3i-3g-3a-Cter quaternary complex on the 40S subunit interface was also observed in a new map we derived for our previous py48S-closed complex by masked classification with signal subtraction around the β-propeller of eIF3b (Llácer et al., 2015). It is now evident, therefore, that the entire eIF3b-3i-3g-3a-Cter module re-locates from the solvent side to the 40S subunit interface of the open conformation of the PIC, and remains there in the closed conformation up to start codon selection, with the eIF3b/Prt1 subunit making multiple contacts at the subunit interface with some alterations between the two states.

The different PIC structures showing the eIF3b-3i-3g-3a-Cter module bound to the solvent surface (py48S-5N) or at its alternative location at the subunit interface (py-48S-eIF3-open and py48S-eIF3-closed), are of sufficiently high resolution to identify particular residues in eIF3b in proximity to residues in rRNA, eIF1 or eIF2γ at the subunit interface, and a different set of eIF3b residues that closely approach rRNA residues on the solvent side of the 40S subunit. Armed with this information, we carried out genetic analyses of substitutions of key eIF3b residues predicted to perturb its contacts at the subunit interface, observed in either the open or closed conformations, or of eIF3b contacts at the solvent side of the 40S subunit visualized in the late-stage PIC with eIF5-NTD replacing eIF1. The results indicate that eIF3b interactions at the subunit interface expected to stabilize the closed complex, as well as other interactions expected to promote relocation of the eIF3b-3i-3g-3a-Cter module back to the 40S solvent side, are critical for efficient utilization of near-cognate start codons in vivo. These findings establish the physiological importance of direct interactions made by the eIF3b-3i-3g-3a-Cter module at the 40S subunit interface, and support the notion that proper movement of the eIF3b-3i-3g-3a-Cter module between its alternative binding sites in the PIC contributes to the fidelity of start codon selection in vivo.

## Results and Discussion

### Reconstitution and overall structure of the 48S PIC in open conformation

In order to trap the yeast 48S PIC in an open conformation, we improved the reconstitution of this complex by including an unstructured capped 49-mer mRNA with an AUC near-cognate start codon (5′GGG[CU]_3_[UC]_4_UAACUAUAAAA**<u>AUC</u>** [UC]_2_UUC[UC]_4_GAU 3′) in place of the 25-mer uncapped mRNA used earlier (Llácer et al., 2015). Moreover, recombinant eIF4 factors were expressed in bacteria and purified, and the reconstituted eIF4F complex and eIF4B were used to activate the mRNA. The resulting eIF4-mRNA complex was then mixed with pre-formed 43S PIC to form 48S PICs. In assembling 43S PICs, we used eIF3 purified from yeast with a modified purification protocol that minimizes co-purification of eIF5 rather than the recombinant eIF3 expressed in bacteria used in our earlier study, which lacked N-terminal segments of eIF3c and eIF3g (Llácer et al., 2015). Cryo-grids were prepared, a large data set of more than 2600 images was collected, and signal subtraction and 3D classification using masks were enlisted during image processing to obtain a 48S PIC in an open conformation at an overall resolution of 5.2 Å (Figures 1A, 1B and Figure 1 – figure supplement 1). This PIC, henceforth termed ‘py48S-open-eIF3’, is similar to that of the py48S-open PIC described previously (Llácer et al., 2015) except for the presence of clear density for the PCI domains of eIF3 at the solvent side, and improved density for eIF3 domains at the 40S subunit interface. py48S-open-eIF3 and the py48S-open complex reported earlier have similar conformations of the 40S subunit and positioning of all non-eIF3 factors, except for a few extra residues of the eIF1 N-terminal tail observed approaching and making contacts with eIF2γ in the new map (Figure 1C). Hence, we shall mainly discuss here the PCI domains of eIF3, not observed earlier, and the eIF3 domains observed at the subunit interface at higher resolution, compared to our previous py48S-open complex.

**Figure 1.**
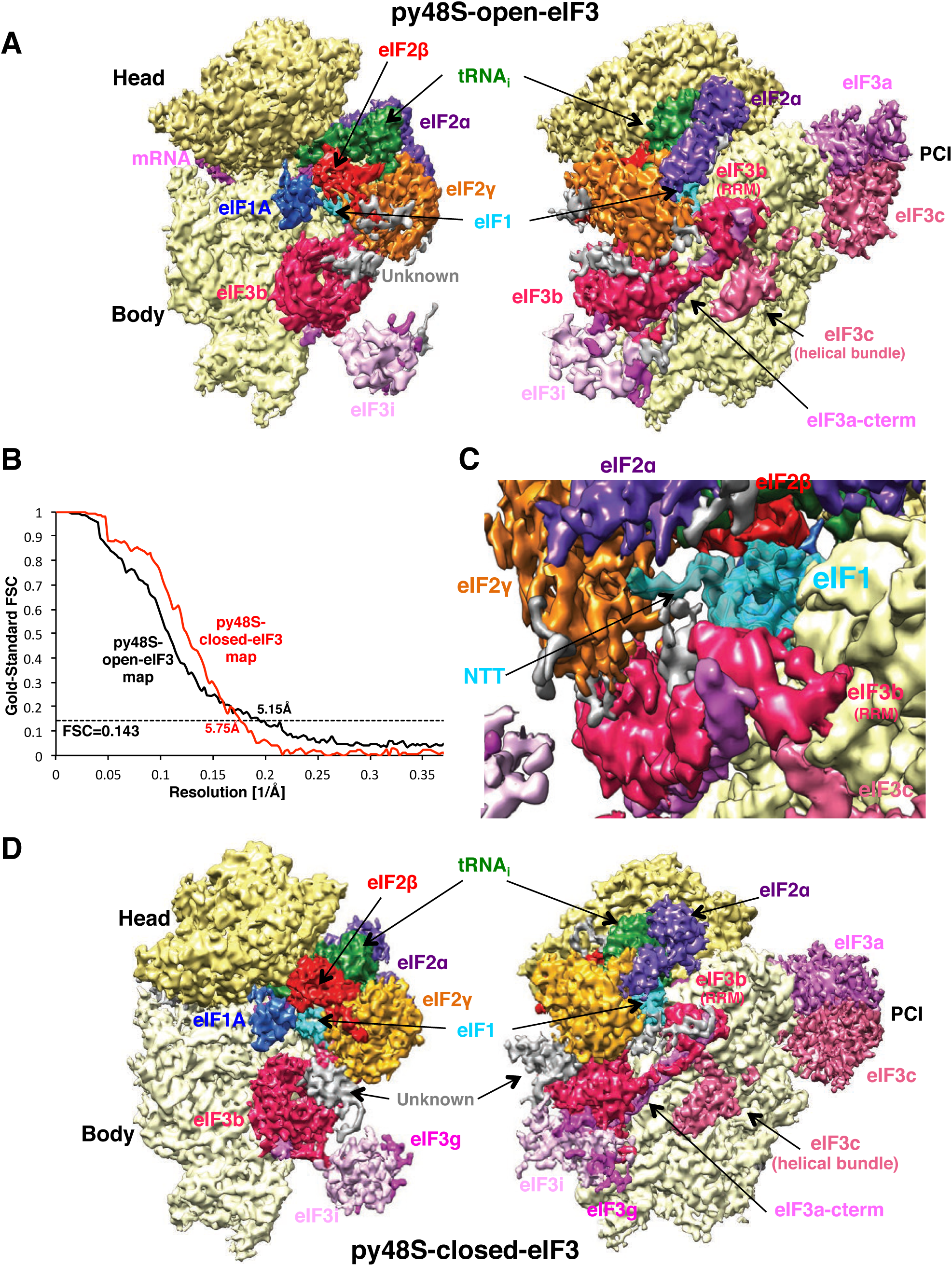
Cryo-EM structure of the py48S-open-eIF3 and py48S-closed-eIF3 PICs. (A) Cryo-EM maps of the p48S-open-eIF3 PIC shown in two orientations. Regions of the map are colored by component to show the 40S subunit (yellow), eIF1A (blue), eIF1 (cyan), Met-tRNA_i_^Met^ (green), mRNA (magenta), eIF2α (violet), eIF2γ (orange), eIF2β (red), eIF3 (different shades of pink). The 40S head is shown in a darker yellow compared to the body. The same colors are used in all the figures. (B) Gold-standard Fourier Shell Correlation (FSC) curves for the py48S-open-eIF3 and py48S-closed-eIF3 maps. (C) eIF1 N-terminal tail (transparent cyan surface) approaching and making contacts with eIF2γ (D) Cryo-EM maps of the py48S-closed-eIF3 PIC shown in two orientations.

The aforementioned newly available image processing tools were also used to reprocess our earlier reported ‘py48S-closed’ PIC (Figure 1 – figure supplement 1) to obtain a map at an overall resolution of 5.75 Å resolution showing clear densities for the different eIF3 domains at the 40S subunit interface, henceforth termed ‘py48S-closed-eIF3’ PIC (Figures 1B & 1D). The resolution of the ‘py48S-closed-eIF3’ PIC map is lower than that of the previously reported ‘py48S-closed’ PIC, probably because of the relatively smaller number of particles used in the current reconstruction (12,732 vs. 21,401 particles). However, the particles used to obtain the new structure seem to be relatively more homogenous for the different eIF3 domains at the subunit interface. Hence, we obtained much improved density (Figure 1 – figure supplement 2) for these eIF3 domains despite overall lower resolution for the whole PIC. The py48S-closed-eIF3 PIC reveals a closed conformation of the 40S subunit, similar to our earlier reported py48S-closed (Llácer et al., 2015) and py48S complexes (Hussain et al., 2014). Moreover, eIF3 is observed in a similar conformation, with the PCI domain heterodimer of eIF3a/-3c positioned at the solvent side and other eIF3 domains located at the 40S interface, compared to both the previous py48S-closed and the new py48S-open-eIF3 PIC. Accordingly, we shall only discuss the densities corresponding to eIF3 domains located at the 40S subunit interface in the py48S-closed-eIF3 map since these are the only parts of the map with better local resolution compared to those in our previously reported py48S-closed map.

### PCI domains of eIF3a and -3c bound on the solvent side of the 40S subunit

The density for the PCI domains of eIF3a and -3c are clearly observed in py48S-open-eIF3 on the solvent side of the 40S (Figures 1A and 2A). The overall positions of these domains are similar to those observed in the py48S-closed-eIF3 complex (Figures 1A, 1D), showing that the PCI domains remain anchored to the solvent interface in both the open and closed states of the 48S PIC. Upon superimposition of the coordinates of the 40S body in py48S-open-eIF3 and py48S-closed-eIF3, we observe a shift in the position of the PCI domains in the two states (Figure 1 – figure supplement 3A), which may be necessary to facilitate the distinct repositioning of other eIF3 domains at the subunit interface in the transition between the two states (discussed later). It is likely that using eIF3 purified from yeast for reconstituting the 48S complex and also applying an eIF3 mask for 3D classification during image processing has helped in isolating a class of particles with higher occupancy of eIF3 in the open state compared to our earlier effort with py48S-open (Llácer et al., 2015).

**Figure 2.**
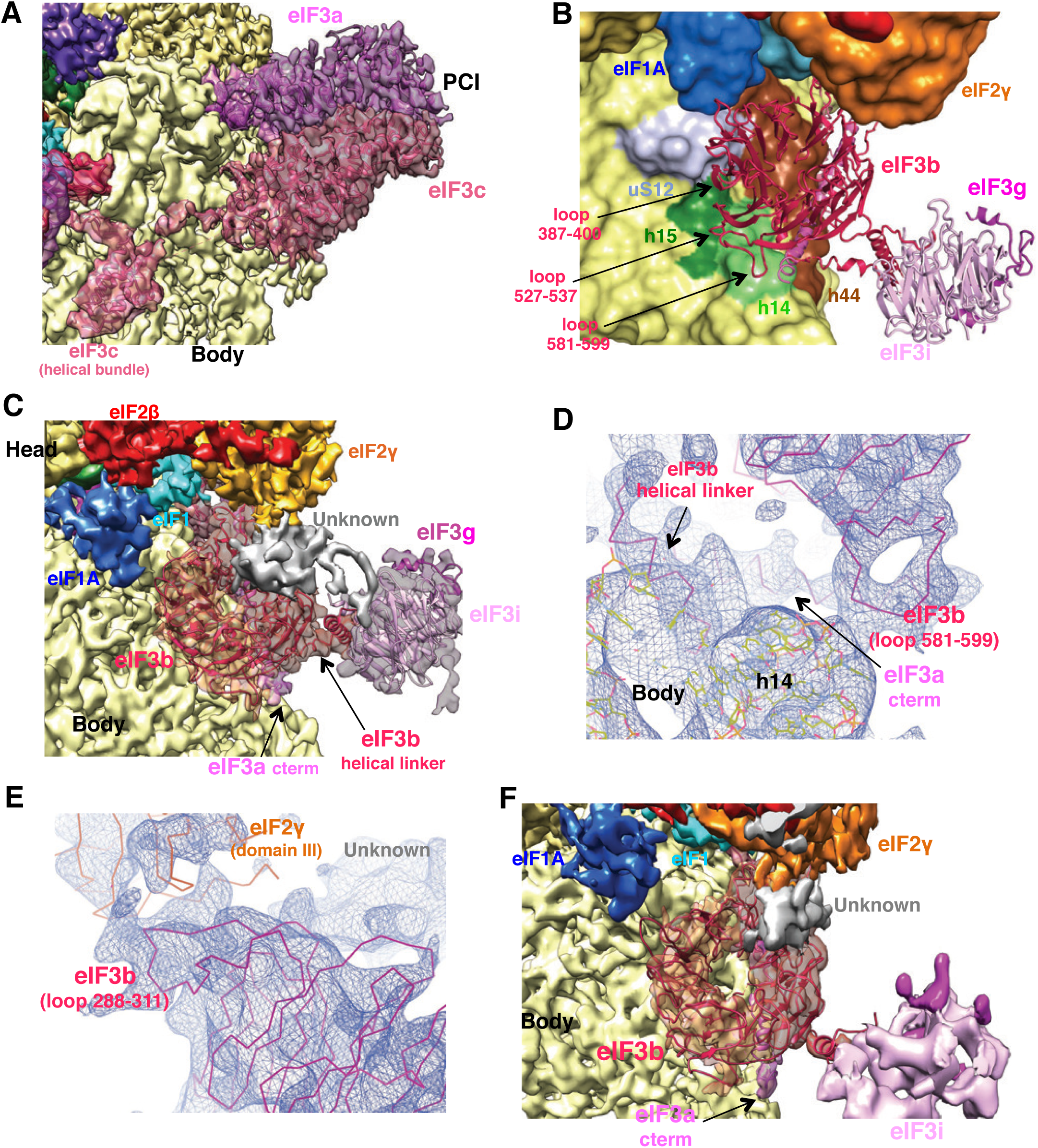
Contacts of the eIF3 PCI heterodimer and of the eIF3b-β propeller with other components in the 48S PIC. (A) Fitting of the eIF3a/c PCI heterodimer in the py48S-open-eIF3 map. (B) View of the contacts of the eIF3b β-propeller with different parts of the 40S subunit (colored variously), eIF2γ and the eIF3a-CTD. (C) Fitting of the eIF3b and eIF3i β-propellers into the py48S-closed-eIF3 map. An unknown density on top of the eIF3b β-propeller and in contact with eIF2γ is shown in gray. (D,E) Fitting of some well-resolved loops of the eIF3b β-propeller into the py48S-closed-eIF3 map. (F) Fitting of the eIF3b and eIF3i β-propellers into the py48S-open-eIF3 map.

### eIF3b bound on the subunit interface of 40S

Compared to the previous open and closed complexes (Llácer et al., 2015), the density for eIF3 domains at the 40S subunit interface is more clearly resolved in the current py48S-open-eIF3 and py48S-closed-eIF3 maps (Figures 1A, 1D and Figure 1 – figure supplement 2). Density for a β-propeller on the 40S subunit interface near h14 and h44 and in contact with eIF2γ was observed, as reported earlier (Llácer et al., 2015) (Figures 1A, 1C, 1D and 2B). Previously, we had tentatively assigned this density to the β-propeller of eIF3i since partial density for the β-propeller of eIF3b could be seen at its well-known location at the 40S solvent interface near h16 (Llácer et al., 2015). However, this density on the 40S subunit interface near h14 and h44 was re-interpreted as the eIF3b β-propeller by docking an improved mammalian model of eIF3b into the yeast ‘py48S-closed’ PIC map (Simonetti et al., 2016). These authors were guided by the observation that the β-propeller of eIF3i appeared to be bound instead at the GTPase-binding site on the subunit interface in the mammalian 48S late stage IC described in their study (Simonetti et al., 2016). However, this density assigned to the eIF3i β-propeller was also later re-interpreted as ABCE1 protein (Heuer et al., 2017; Valášek et al., 2017) (Mancera-Martínez et al., 2017).

In view of these previous difficulties in identifying the β-propellers of eIF3i and eIF3b in low-resolution maps, it was important to be certain about their assignments in our current models. With our improved maps for the β-propeller bound near h14 and h44 on the 40S subunit interface, we can unambiguously assign this density to the β-propeller of eIF3b, as it shows a decidedly better fit to the density compared to that of eIF3i obtained previously (Llácer et al., 2015). The diameter of the inner and outer rims of the ring-shaped density are more appropriate for the size of the 9-bladed β-propeller of eIF3b (Figures 2C and F) versus the 7-bladed β-propeller of eIF3i. Furthermore, density corresponding to loops present in the eIF3b and not in the eIF3i β-propeller are also observed (Figures 2D and 2E). Thus, we agree with Simonetti et al. (Simonetti et al., 2016) that the β-propeller of eIF3b instead of the β-propeller of eIF3i is bound near h14 and h44 in the previous py48S-closed map, and we show this here as well in the maps of both py48S-open-eIF3 and py48S-closed-eIF3. However, no density was observed at the GTPase-binding site of the 40S for the β-propeller of eIF3i (see below for eIF3i’s position). Also, in these maps we do not observe density for the β-propeller of eIF3b at the solvent interface near h16 (Figure 1 – figure supplement 3B). It is likely that our previously reported py48S-open and py48S-closed maps (Llácer et al., 2015) had a small fraction of 48S PIC particles with the eIF3b β-propeller in its alternative position at the solvent interface (Hashem et al., 2013) (Aylett et al., 2015) (Llácer et al., 2018) (des Georges et al., 2015) (Eliseev et al., 2018) in addition to those harbouring this eIF3b domain at the subunit interface.

The eIF3b β-propeller is anchored between h15, h44 and ribosomal protein uS12 using a loop (residues 387-400) and other two loops in the β-propeller (residues 527-537 and 581-599) are also in contact with h14 (Figures 2B, 2D). Although the β-propeller is positioned in proximity of eIF1A, no direct interaction between the two was observed. Another eIF3b loop attached to the β-propeller (residues 288 to 311) is observed in contact with domain III of eIF2γ in both py48S-open-eIF3 and py48S-closed-eIF3 PICs (Figure 2E). The eIF3b RRM (RNA recognition motif) domain is also observed in py48S-open-eIF3 and py48S-closed-eIF3 PICs (Figures 3A, 3B) interacting with h24 at the 40S platform, with a long helix in the eIF3a C-terminal region (the same long helix is also in contact with the β-propeller of eIF3b), and with eIF1 and the eIF3c N-terminal tail (see below) (Figure 3C). Interaction of these eIF3 elements with the 40S or the other initiation factors eIF1 and eIF2 promotes a distinctly different conformation of the eIF3b/eIF3a-CTD subcomplex at the subunit interface compared to the same subcomplex bound at the 40S solvent interface (Figure 3D).

**Figure 3.**
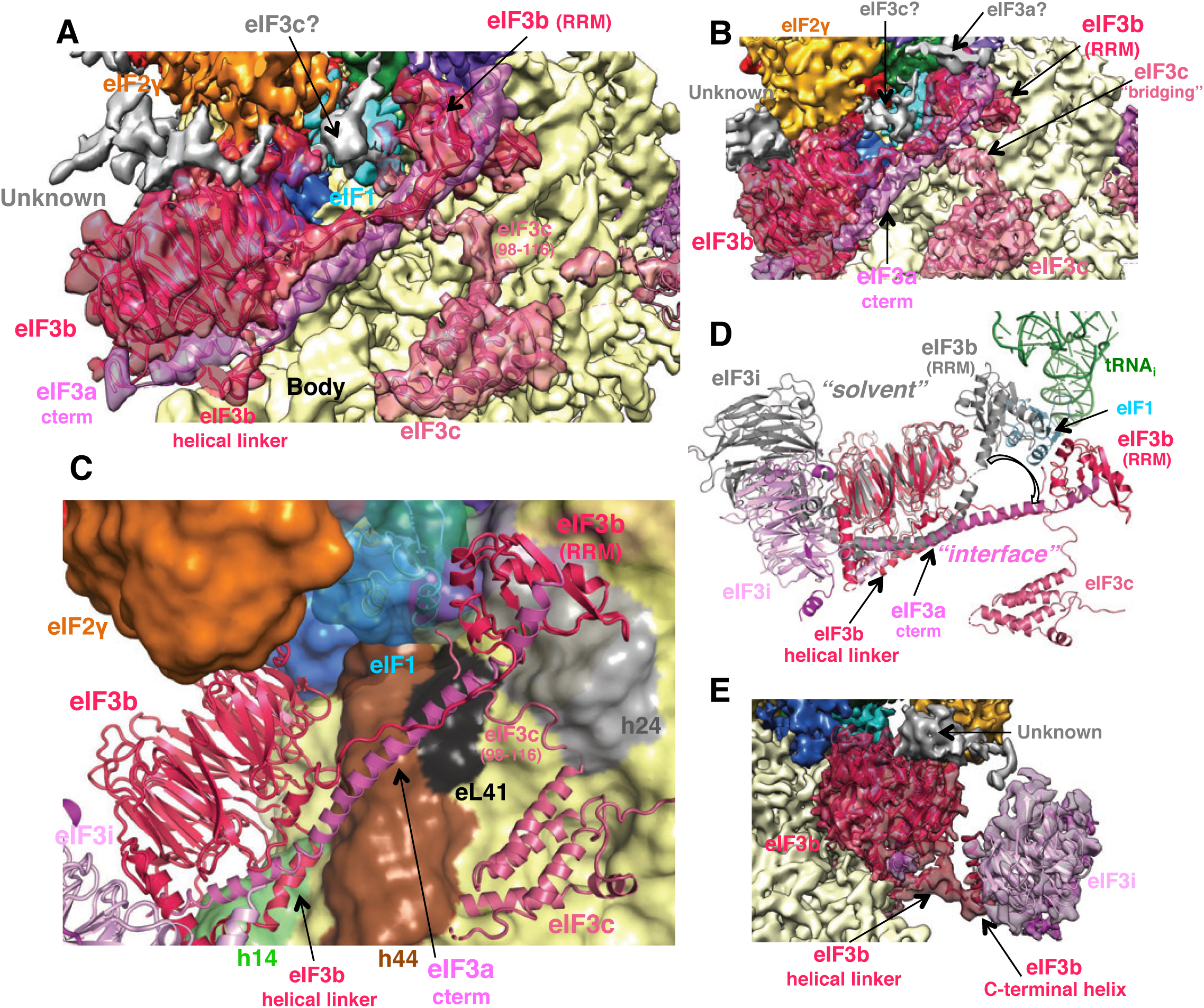
Contacts of eIF3b, the eIF3a C-term and the eIF3c N-term with other components in the 48S PIC. (A) Fitting of all of eIF3b, the eIF3a C-term and the eIF3c N-term into the py48S-open-eIF3 map. (B) As in A, fitting of all of eIF3b, the eIF3a C-term and the eIF3c N-term into the py48S-closed-eIF3 map. (C) View of the contacts of the eIF3b β-propeller, eIF3b RRM domain, eIF3a-CTD and most of the N-terminal portion of eIF3c with one another and with different parts of the 40S subunit (colored variously), eIF2γ and eIF1. (D) Superimposition of the eIF3b/eIF3i/eIF3g/eIF3a-Cterm quaternary complex observed in py48S-closed-eIF3 with that found on the 40S solvent side in py48S-eIF5N (in grey; PDB: 6FYY), aligning the eIF3b β-propellers, shows how this eIF3 subcomplex undergoes internal rearrangements in the transition between the two states, possibly resulting from constraints imposed by its interactions with eIF2 and eIF1 unique to the subunit interface location. (E) Fitting of the eIF3b and eIF3i β-propellers in the py48S-closed-eIF3 map in a different view from that in Figure 3C to highlight the fitting of the eIF3b helical linker connecting the two different eIF3 β-propellers.

The possible interactions of eIF3b residues with residues in rRNA, 40S proteins, or other eIFs at the subunit interface in the open and closed conformations are listed in Table 1 and Figure 3 – figure supplement 1. Although the eIF3b β-propeller is positioned similarly between h14 and h44 in both py48S-open-eIF3 and py48S-closed-eIF3 PICs, a subtle repositioning of the β-propeller in the closed state leads to some differences in its interactions with 40S or eIF residues between the two states (Movie 1). Some residues of eIF3b are predicted to make the same interactions with the PIC in both states, whereas a few residues engage in a certain interaction only in the open or closed complex (Table 1; see also genetic studies below), including direct contact between the eIF3b β-propeller and eIF1 observed exclusively in the open state (Table 1). The latter interactions should help keep eIF1 positioned at the P site during scanning, while the lack of eIF3b/eIF1 interaction in the closed state would favour eIF1 release from the PIC after recognition of the start codon. Interestingly, in the closed state only, several acidic side chains of the eIF3b β-propeller (Glu308, Asp397, Glu398) are positioned in proximity to the negatively charged phosphate residues of rRNA and acidic side chains of eIF2γ, consistent with electrostatic repulsion. This repulsion might help to promote the re-positioning of eIF3b back to the solvent interface after eIF1 leaves the PICs. Similar repulsion was predicted between eIF1 and Met-tRNA_i_ in the closed-state PICs where acidic side chains of eIF1 are positioned in proximity to the negatively charged phosphate residues of tRNAi (Hussain et al., 2014) and shown to promote initiation accuracy in vivo (Thakur and Hinnebusch, 2018); and similarly between IF3 and fMet-tRNA_i_ in the closed state of bacterial PICs (Hussain et al., 2016).

**Table 1:** Possible interactions of eIF3b residues with the 40S subunit or other eIFs in the py48S-open-eIF3 or py48S-closed-eIF3 complexes*

| eIF3b residue | Interacting residue(s) in py48S-open-eIF3 | Interacting residue(s) in py48S-closed-eIF3 |
| --- | --- | --- |
| P88 | rRNA 986G | same |
| K91 | rRNA 1011U | rRNA 1011U, 1012A and 986G |
| S101 | rRNA 995U | rRNA 994A |
| K105 | R53 and D50 of eIF1, D97 and phosphoSer98 of eIF3c | None |
| T120 | None | rRNA 984G, 985G |
| K122<sup>#</sup> | rRNA 986G and 985G | rRNA 985G |
| K142 | D50 and Y49 of eIF1, NTD of eIF3c | None |
| S143 | R53 of eIF1, Y96 of eIF3c | None |
| K147<sup>#</sup> | Y96 and phosphoSer99 of eIF3c | same |
| R148 | S98, phosphoSer 99, D100 and phosphoSer103 of eIF3c | phosphoSer 99, D100 and phosphoSer103 of eIF3c |
| D150 | rRNA 1011U | same |
| K152<sup>#</sup> | rRNA 982A and 1012A | rRNA 981U and 982A |
| Q250 | K252 of eIF2γ | R454 and K252 of eIF2γ |
| Y268 | R454 of eIF2γ | same |
| D271 | None | K252 of eIF2γ |
| E290 | R439 of eIF2γ | unknown density on top of eIF3b propeller (eIF3g?, eIF5 CTD?) |
| I293 | K452 of eIF2γ | I505 and E506 of eIF2γ |
| V294 | A451 of eIF2γ | I503 of eIF2γ |
| E295 | R454 of eIF2γ | R454 of eIF2γ |
| E296 | None | R504 and K207 of eIF2γ |
| D297<sup>#</sup> | D457 and R454 of eIF2γ | D460 of eIF2γ |
| E308<sup>#</sup> | Residues around 445-448 | E506 of eIF2γ (may be with K449 of eIF2γ as well) |
| K346<sup>#</sup> | rRNA 1742A | None |
| K363 | rRNA 1743G and 1744A | rRNA 1745G |
| M366 | rRNA 1745G | None |
| F393 | Q75 of uS12 | None |
| R394 | rRNA 433G and I77 of uS12 | Q75 and L76 of uS12 |
| N395 | rRNA 46A | rRNA 433G |
| G396 | None | rRNA 1742A |
| D397<sup>#</sup> | None | rRNA 1743G |
| E398<sup>#</sup> | None | rRNA 1743G |
| R424 | Q75 of uS12 | L54 and E55 of uS12 |
| R426 | E 98 and N99 of uS12 | N99 of uS12 |
| V427 | None | N99 of uS12 |
| R476 | rRNA 431G | E101 of uS12 |
| D477 | None | S145 of uS12 |
| E530 | rRNA 413C | None |
| K531 | rRNA 413C and 415A | rRNA 413C |
| T532 | rRNA 412U and 413C | None |

\* At the current resolution, side chains of residues are not well resolved, and therefore this table reflects possible interactions with residues in close proximity to a given eIF3b residue. As a consequence, in some cases we include more than one possible interacting residue. Possible contacts of eIF3b residues located in the helical linker between the β-propeller and the C-terminal-eIF3i-interacting helix (residues 668-701) with the 40S and eIF3a-cterm are not mentioned here because we have modeled “de novo” only the backbone but not the side chains for residues on this region.
\*\* Interactions of eIF3b with eIF3a CTD are not mentioned as they are largely similar in the OPEN and CLOSED states and may involve residues Y78, V80, N82, E113, F126, F128, Y158, V163, Y166, N167, N170, D172, T173, F175, E177, P178, P181, T182, P185, S187, K190, L193, M194, V198, R559, F560, D580, Y583, P584, G585, K600, V602, R620, I642, A643, R663, residues of the helical linker (see above).
\*\*\* Interactions of eIF3b with 3i are not mentioned because of the poor local resolution of this part of the map. In any case, we do not expect differences in the contacts between eIF3i and eIF3b to those observed in the high resolution crystal structures of eIF3i-eIF3b yeast complexes, PDB: 3ZWL or PDB: 4U1E.

### Trimeric complex of eIF3i-3g-3b C-terminal helix bound to the eIF3b β-propeller also re-locates to the subunit interface of 40S

Most of the residues in the linker region between the C-terminal helix of eIF3b (in contact with eIF3g) and the β-propeller of eIF3b are observed in py48S-open-eIF3 and py48S-closed-eIF3 PICs maps (Figures 2C, 2D, 2F, 3A and 3E). This eIF3b helical linker region is in contact with h13 and h14 of rRNA (Figure 2D and 3C) and with the previously mentioned C-terminal helix of eIF3a (Figure 3A, 3B and 3C). These interactions between the eIF3b helical linker region and C-terminal helix of eIF3a are similar to those occurring among the same eIF3 regions observed at the solvent side of the 40S in both mammalian 43S (des Georges et al., 2015; Valášek et al., 2017) and yeast py48S-5N maps (Figure 3D) (Llácer et al., 2018). Furthermore, the C-terminal helix of eIF3b was earlier shown to interact with the β-propeller of eIF3i (Erzberger et al., 2014) (Herrmannová et al., 2012; des Georges et al., 2015; Valášek et al., 2017) (Figure 2C and 3E). Together, these findings suggest that the interactions between the two β-propellers of eIF3b and eIF3i may be intact at the solvent surface as well. Interestingly, at a lower resolution threshold of the maps, density for the trimeric complex eIF3i-eIF3g-eIF3b C-terminal helix can be observed in close proximity to the eIF3b β-propeller in both py48S-open-eIF3 and py48S-closed-eIF3 PICs (Figures 2C and 3E). Thus, it appears that both β-propellers of eIF3b and eIF3i and the regions of eIF3g interacting with eIF3i move together as a module to the subunit interface from the solvent surface. Also, the β-propeller of eIF3i appears to be flexible and not restricted to one conformation, resulting in weak density, and it does not contact the 40S. Finally, there is an unassigned density on top of the β-propeller of eIF3b in contact with eIF2γ, which is more prominently seen in py48S-closed-eIF3 (Figures 1A, 1D, 2C, 2F, 3A, 3B, 3E and Figure 1 – figure supplement 2B), which might correspond either to the RRM domain of eIF3g based on its size and proximity to eIF3i or to the C-terminal domain of eIF5 based on the contacts of eIF2γ with the structurally homologous heat domain of eIF2Bε (Kashiwagi et al., 2019; Kenner et al., 2019); although further studies are required to identify it with confidence.

### N-terminal region of eIF3c bound on the subunit interface of 40S

Clear density is observed for different parts of the N-terminal region of eIF3c in py48S-open-eIF3 and py48S-closed-eIF3 PICs. In our earlier studies we had observed a helical bundle of eIF3c on the subunit interface near h11/h24/h27 (Llácer et al., 2015). In the new py48S-open-eIF3 map, connectivity between the 5 helices can also be seen. Using our previous py48S-closed map (at slightly higher overall resolution, 4.9 Å), the newer py48S-open-eIF3 and py48S-closed-eIF3 maps, and secondary structure predictions (Figure 4A), enabled us to model the N-terminal region of eIF3c (residues 117 to 226) in both py48S-open-eIF3 and py48S-closed-eIF3 maps, including certain amino acid side chains that interact with rRNA side chains (Figure 4B, 4C and 4D). In our proposed model for the eIF3c helices, most of the hydrophobic residues are buried and therefore protected from the solvent (Figure 4E), whereas charged residues are exposed and may interact with rRNA, eIF1, eIF3b RRM, or other ribosomal proteins (Figure 4E and 4F). Also, a clear eIF3c density (residues 98-116) connecting the eIF3c helical bundle to eIF1 and the eIF3b RRM is observed (Figures 3A, 3B, 3C, 4E and 4F), bridging eIF1 and the eIF3b RRM. This might explain why, unlike the well-established eIF1:eIF3c-NTD interaction, a direct interaction of eIF1 with the eIF3b RRM has never been reported, as it might need concurrent interaction of eIF1 with a fragment of eIF3c to be stabilized. This putative “bridging” eIF3c density approaches the residues of eIF1 (50-56) that were earlier proposed to interact with eIF3c (Reibarkh et al., 2008). The basic sidechains of eIF3c residues 109 to 116 interact with h24 of the rRNA, while residues 98-107, which are acidic in nature, interact with basic residues of eIF1 (residues K52, R53, K56, see above), the RRM of eIF3b (around residues 145-154) and eL41 residues 19-23 (Figure 4E and 4F). Most of the observed eIF3c residues involved in these contacts are highly conserved (Figure 4A). Additional density attached to eIF1 is observed in both py48S-open-eIF3 and py48S-closed-eIF3 maps and may correspond to the remaining N-terminal residues (1-96) of eIF3c engaged in additional interactions with eIF1 (Fig. 3A-B, grey density). However, further studies with better maps will be required to confirm this last possibility..

**Figure 4.**
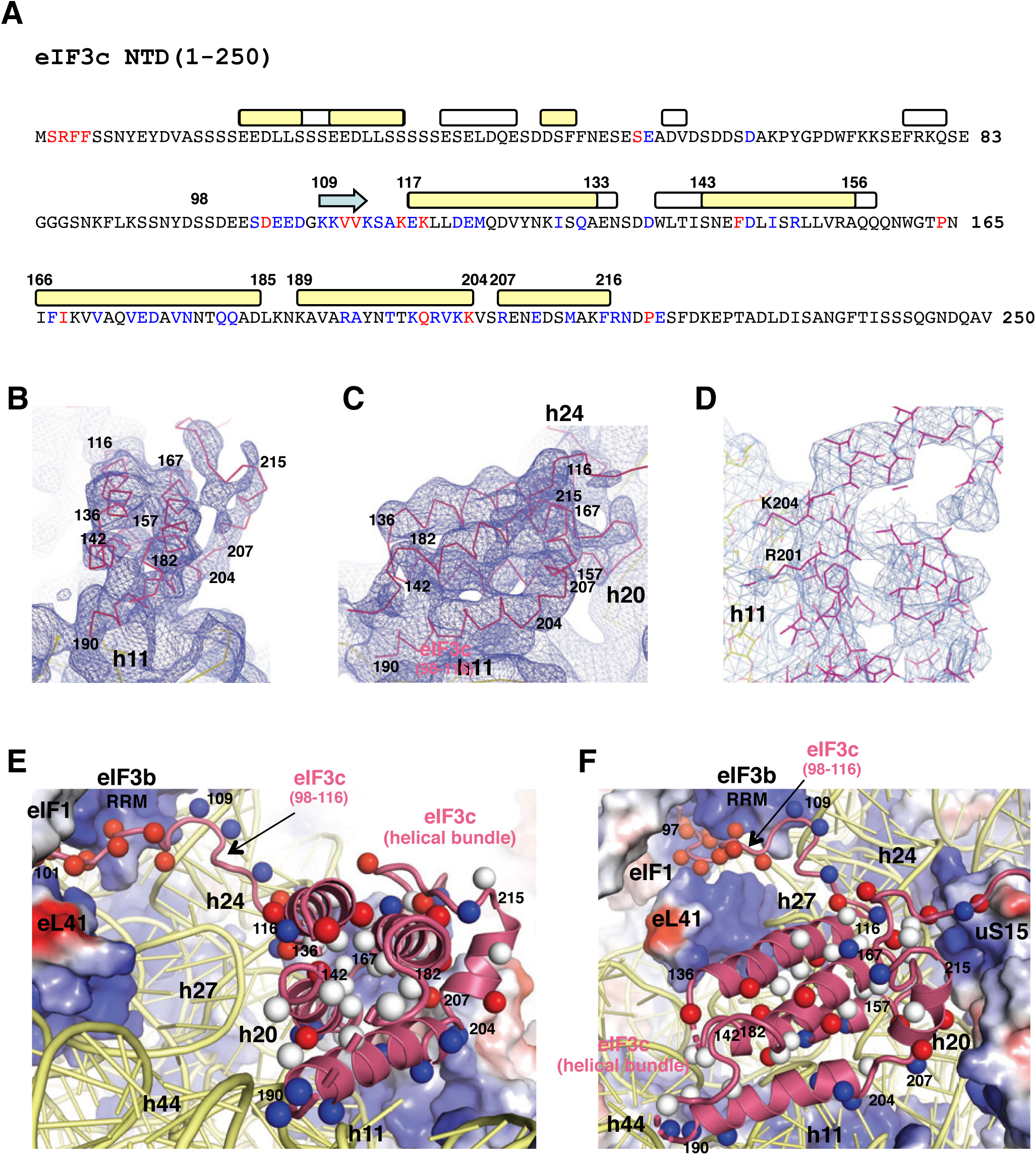
Proposed model for the N-terminal region of eIF3c and its contacts with other components in the 48S PIC. (A) Amino acid sequence, secondary structure prediction and sequence conservation of the eIF3c N-terminal region. The β-strands and α-helices are depicted as arrows and rectangles, respectively. Yellow rectangles account for the consensus between the three programs used for secondary structure prediction (see Methods section), whereas those in white correspond to the prediction of at least two of these programs. Residues shown in red are invariant whereas residues with only conservative replacements are in blue in a multiple sequence alignment (see Methods section). (B,C) Two different views of the eIF3c 5-helical bundle showing the connectivity between the five helices. Residue numbers at the beginning and end of each helix are labeled, exhibiting substantial agreement with the secondary structure predictions in (A). (D) The side chains of several residues in the eIF3c 5-helical bundle are visible, especially those in contact with rRNA. Two very conserved basic residues (R201, K204) interacting with h11 are labeled. (E,F) Surface electrostatic representations of 40S proteins, eIF1, and the eIF3b RRM domain and cartoon representations of rRNA and the eIF3c N-terminal region. The C_β_ for each residue with hydrophobic, basic or acidic sidechains of eIF3c is represented as white, blue and red spheres, respectively. Residue numbers at the beginning and end of each helix are also labeled. In (E), it can be seen that most of the hydrophobic residues are buried, whereas in (E) and (F) the basic residues are exposed and interact mainly with rRNA, whereas the acidic residues interact with the basic surfaces of eIF1 and eIF3b RRM, as well as with eL41 and uS15.

### C-terminal region of eIF3a bound on the subunit interface of 40S

A long helix of ∼80 residues of the eIF3a C-terminal region is observed running alongside the β-propeller of eIF3b and interacting with both the face of the eIF3b RRM β-sheet and the eIF3b linker connecting the RRM to the β-propeller, thereby providing an additional connection between these two eIF3b domains, in both the py48S-open-eIF3 and py48S-closed-eIF3 maps (Figure 3A, 3B and 3C). The most N-terminal residues of this eIF3a helix run underneath the eIF3b β-propeller (Figures 3A and 3B) and also make contacts with h14 and h44 of the 40S (Figure 3C) and with the eIF3b helical linker that connects the eIF3b β-propeller to the eIF3b C-terminal helix that interacts with eIF3i (Figures 3A-C) (see above). The density is not clear enough, however, to define unambiguously the side-chains of this eIF3a helix. A completely stretched linker of at least 110 amino acids would be needed to connect the eIF3a helix to the eIF3a PCI domain at the solvent side, a distance of more than 300 Å around the 40S (Figure 5A-C). Based on secondary structure predictions, sequence conservation among homologs, known biochemical information, and these distance constraints, we have tentatively assigned this linker to the region at the eIF3a C-terminal end spanning residues 693-771. Mutations in this region of eIF3a, including those for the KERR motif (Chiu et al., 2010; Khoshnevis et al., 2014) and/or substituting residues 692-701 with Ala (Szamecz et al., 2008), are known to impair the binding of eIF3a to the RRM of eIF3b and to the 40S ribosome, respectively. Finally, there is an unassigned density at the extreme C-terminus of the eIF3a helix that also approaches eIF2γ, which probably belongs to eIF3a, but the low resolution of this density and the fact that there are still some segments of factors not accounted for, prevents us from assigning this density unambiguously (Figure 3B).

**Figure 5.**
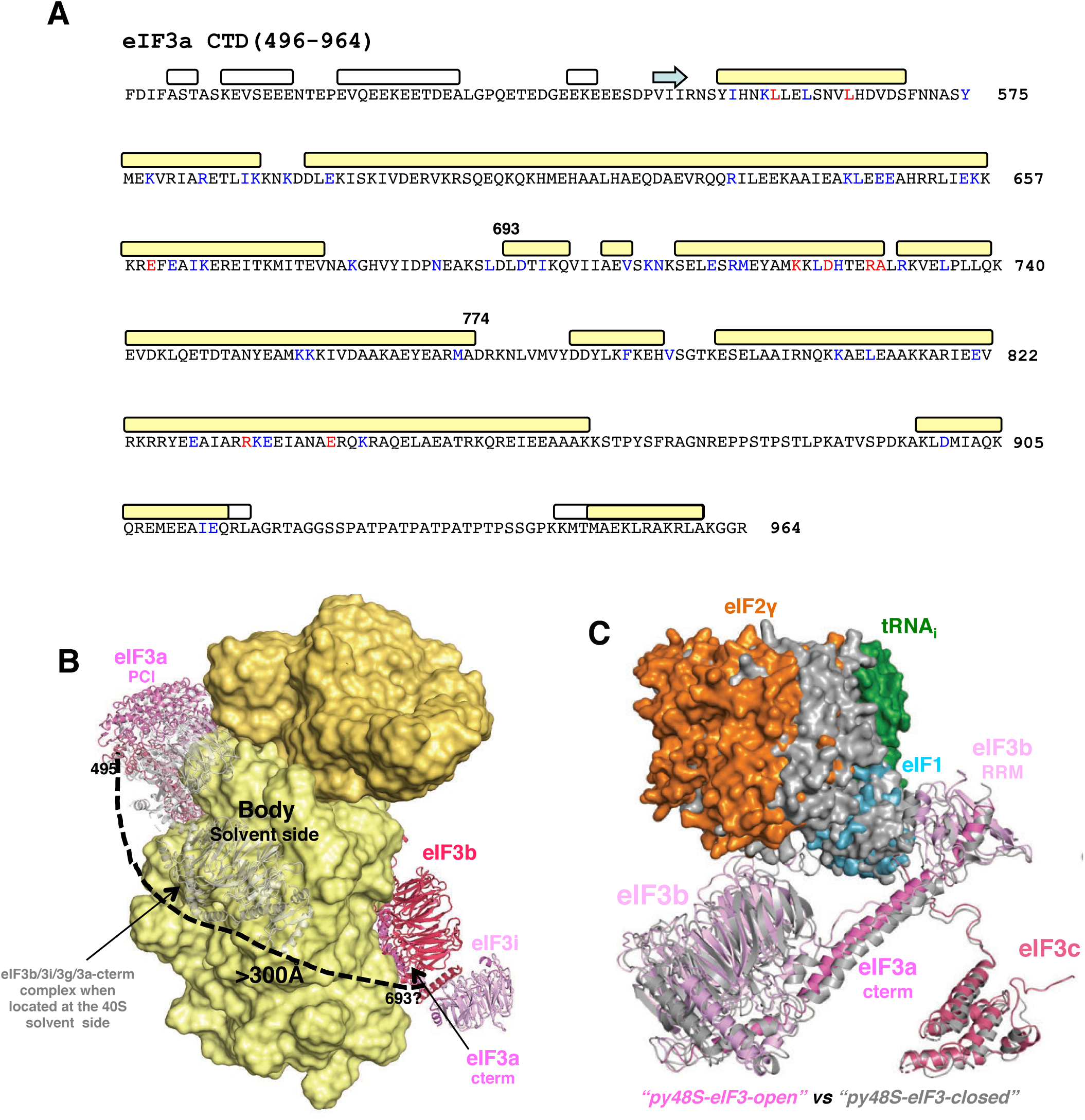
Modeling of the eIF3a C-terminal helix. (A) Amino acid sequence, secondary structure prediction and sequence conservation of the eIF3a C-terminal region, all depicted as in Figure 4A. (B) Proposed path along the solvent side of the 40S subunit for the central portion of eIF3a connecting the eIF3a/PCI domain and eIF3a C-term helix in py48S-closed-eIF3. A linker more than 300 Å long is needed to connect the C-end of the eIF3a PCI domain with the N-end of the eIF3a C-terminal helix that interacts with eIF3b, spanning eIF3a residues 495-693. A semitransparent cartoon representation of the eIF3b-3i-3g-3a-Cter complex is shown at the subunit interface connected to the predicted central eIF3a linker on the solvent-exposed side of the 40S subunit (dashed line), which is presumed to be fully extended to permit re-location of the eIF3b-3i-3g-3a-Cter module to the 40S subunit interface. (C) Superimposition of eIF3 subunits in the py48S-closed-eIF3 and py48S-open-eIF3 structures, achieved by aligning the 40S bodies in both structures. For clarity, the eIF3i/eiF3g subunits are not shown. tRNA_i_, eIF2 and eIF1 are shown as surfaces. Other components of the py48S-closed-eIF3 structure are colored gray.

### mRNA path in the py48S-open-eIF3

Since a longer mRNA was used to reconstitute py48S-open-eIF3 compared to the previous py48S-open PIC, mRNA density is now observed at the openings of both the entry and exit portions of the mRNA-binding channel on the 40S in py48S-open-eIF3 (Figure 1A, Figure 1 – figure supplement 3B). However, there is no distinct high-resolution mRNA density throughout the channel outside of the P site. A stretch of 3 nucleotides of mRNA have been modelled with confidence at the P site (Figure 1 – figure supplement 3C), compared to only 2 nucleotides in the previous py48S-open complex (Llácer et al., 2015). These include the AU at positions 1 and 2 of the AUC start codon, observed interacting with the UA of the anticodon of the tRNA_i_, and the A at the -1 position of the mRNA (Figure 1 – figure supplement 3C). There is no distinct density for the C at position 3 of the AUC codon, probably due to the mismatch in codon:anticodon base-pairing. Thus, overall the mRNA seems to have minimal interaction with the widened mRNA channel of the 40S subunit in this open conformation of the 48S PIC, consistent with a state conducive to mRNA scanning.

### eIF3b interactions at the interface of the 48S PIC modulate the fidelity of start codon selection in vivo

The RRM domain of eIF3b/Prt1 contacts eIF1 and the eIF3c-NTD, and the β-propeller of eIF3b contacts eIF2γ, at the 40S subunit interface of both the py48S-open-eIF3 and py48S-closed-eIF3 complexes (Figure 1A, 1C-D, and Figure 3 – figure supplement 1). It is possible that these interactions stabilize the open conformation and promote scanning, eg. by helping to anchor eIF1 on the 40S platform as mentioned above. Alternatively, these interactions might be more important for stabilizing the closed conformation from which eIF1 will be released on start codon recognition, followed by relocation of the eIF3b-3i-3g-3a-Cter module from the subunit interface back to the solvent-exposed surface of the 40S subunit. Presumably, this relocation is necessary to allow eIF5-NTD to occupy the position of eIF1, dissociation of eIF2 from Met-tRNA_i_ and subsequent joining of the 60S subunit (Llácer et al., 2018). We reasoned that if particular eIF3b contacts at the subunit interface are relatively more important for stabilizing the open, scanning conformation, then substitutions perturbing these contacts would shift the system to the closed state and allow inappropriate selection of near-cognate UUG codons in vivo, conferring the “Sui^-^“ phenotype. If instead the contacts preferentially stabilize the closed conformation, or impede the transition from open to closed states, then disrupting them should shift the equilibrium to the open scanning conformation and suppress initiation at UUG codons, for the “Ssu^-^“ phenotype (reviewed in (Hinnebusch, 2014; Hinnebusch, 2017). The Ssu^-^ phenotype might also be conferred by substitutions that impede relocation of the eIF3b-3i-3g-3a-Cter module from the subunit interface back to the solvent side of the 40S, which could result either from strengthening eIF3b contacts on the interface surface or by weakening its contacts on the solvent surface. As summarized below, we identified numerous substitutions predicted to perturb eIF3b interactions on either the subunit interface or the solvent surface of the 40S that confer pronounced Ssu^-^ phenotypes, and a smaller number that produced relatively weak Sui^-^ phenotypes.

To examine the phenotypes of eIF3b substitutions in vivo, we generated the appropriate mutations in the *PRT1* gene on a single-copy plasmid, and introduced the resulting mutant or WT plasmids into a yeast strain in which expression of the chromosomal *PRT1* allele is under control of the galactose-inducible, glucose-repressible *GAL1* promoter. The resulting transformants, isolated on galactose medium, were tested for mutant phenotypes on glucose medium in which expression of the WT chromosomal *PRT1* allele is transcriptionally repressed. The elevated utilization of UUG start codons in Sui^-^ mutants was identified by testing for suppression of the inability of a *his4-303* strain to grow on medium lacking histidine, as *his4-303* eliminates the AUG start codon, and Sui^-^ mutations enable initiation at the third in-frame UUG codon of *HIS4*. Assaying matched *HIS4-lacZ* reporters with a UUG or AUG start codon yields the UUG:AUG initiation ratio, which is elevated by Sui^-^ mutations. Ssu^-^ phenotypes are scored by testing for suppression of both the His^+^ phenotype and elevated *HIS4-lacZ* UUG:AUG initiation ratio conferred in a *his4-303* strain by *SUI5,* a dominant Sui^-^ allele of *TIF5* encoding eIF5-G31R (Huang et al., 1997). Ssu^-^ mutations also generally suppress the inability of *SUI5* strains to grow at 37°C (Martin-Marcos et al., 2014), which we also scored in all of the mutants.

#### (i) Evidence that interactions of eIF3b RRM residues Lys-147 and Lys-152 with eIF3c-NTD and rRNA on the 40S interface preferentially stabilize the closed conformation of the 48S PIC

Lysine-147 in the eIF3b RRM appears to contact the eIF3c-NTD (Fig. 3-figure Supplement 1A) in the vicinity of Tyr-96 and phosphorylated Ser-99 in both py48S-open-eIF3 and py48S-closed-eIF3 (Table 1). Substituting this Prt1 residue with alanine (K147A) confers a strong Ssu^-^ phenotype, suppressing the Slg^-^ at 37°C, His^+^ phenotype at 30°C, and the elevated UUG:AUG *HIS4-lacZ* initiation ratio conferred by *SUI5* (Figure 6A, row 3 vs. row 1, and Figure 6B, cols. 1 & 3) (See Table S1 for a summary of phenotypes of all *prt1* alleles described in this study.) The same three phenotypes characteristic of strong Ssu^-^ mutations were additionally observed for glutamate substitution of Lys-152 (K152E) (Figure 6A, rows 1 & 4; Figure 6B, cols. 1 & 4), which is also in the RRM and appears to contact rRNA helix 24 (Fig. 3-figure Supplement 1A), including residues A982 and A1012 in the open complex and U981/A982 in the closed complex (Table 1). Interestingly, substituting K152 with Ala strongly suppressed the Slg^-^, but only partially reversed the His^+^ phenotype of *SUI5,* and produced a smaller reduction in the UUG:AUG initiation ratio in *SUI5* cells compared to the K152E substitution (Figure 6A, rows 1 & 5; Figure 6B, cols. 1 & 5 vs. 4). The weaker Ssu^-^ phenotype of K152A compared to K152E is consistent with the idea that the Glu substitution introduces electrostatic repulsion with the rRNA phosphate backbone, whereas the Ala substitution would merely eliminates the electrostatic attraction between the Lys side-chain of K147 and the rRNA backbone. None of these substitutions affects cell growth in strains lacking *SUI5* (Figure 6 – figure supplement 1A, rows 6-8 vs. 1), which is typical of Ssu^-^ substitutions in eIF1 (Martin-Marcos et al., 2011).

**Figure 6.**
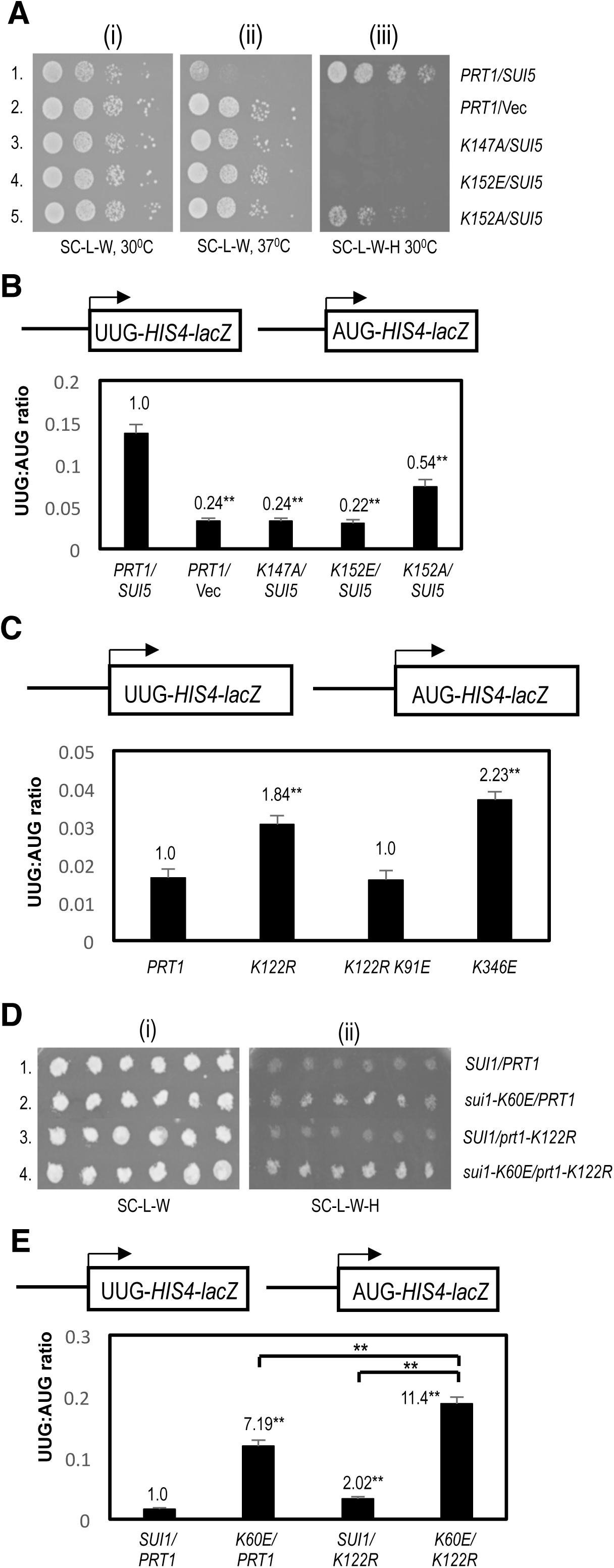
Effect of substitutions in the eIF3b RRM on the fidelity of start codon selection in vivo. (A)-(B) Genetic evidence that weakening interactions of eIF3b RRM residues Lys147 and Lys152 at the 40S subunit interface preferentially destabilize the closed conformation of the 48S PIC and increases discrimination against start UUG codons. **(A)** Serial dilutions of transformants of *P_GAL1_-PRT1 his4-301* strain HD3607 with the indicated *PRT1* alleles on low-copy (lc) *LEU2* plasmids and either single-copy (sc) *TRP1 SUI5* plasmid p4281 or empty vector YCplac22 (Vec) were spotted on synthetic complete medium lacking Leu and Trp (SC-L-W) or the same medium also lacking His (SC-L-W-H) and incubated at 30°C or 37°C for 3-5d. **(B)** The same strains as in (A) harboring *HIS4-lacZ* fusions with AUG or UUG start codons (shown schematically) on plasmids p367 and p391, respectively, were cultured in synthetic minimal medium containing His (SD+His) to an OD_600_ of 1.0-1.2 and β-galactosidase activities were measured in whole-cell extracts. Ratio of mean expression of the UUG and AUG reporters from six transformants are plotted with error bars indicating SEMs. Asterisks indicate significant differences between mutant and WT as judged by a Student’s t-test (*p<0.05, **p<0.01). **(C)-(E) Evidence that weakening interactions of RRM residue Lys122 at the 40S subunit interface preferentially destabilizes the open conformation of the PIC and elevates initiation at UUG start codons. (C)** Transformants of strain HD3607 with the indicated *PRT1* alleles and *HIS4-lacZ* fusions with AUG or UUG start codons were cultured in SD+His+Trp, and analyzed exactly as in (B). **(D)** Transformants of *his4-301* strains HD4108 (*SUI1*) or HD4109 (*sui1-K60E*) with the indicated *PRT1* alleles on lc *TRP1* plasmids were replica-plated to SC-L-W or SC-L-W-H and incubated at 30°C for 3-6d. (E) *HIS4-lacZ* UUG:AUG ratios were determined for strains in (D) exactly as in (B).

Because both K147 and K152 appears to make contacts at the subunit interface in both the open and closed conformations of the py48S PIC, it was possible that substituting these residues would confer dual Sui- and Ssu-phenotypes, as has been observed previously for certain substitutions in eIF5 (Saini et al., 2014). At odds with this possibility, however, neither *K152E* (Figure 6 – figure supplement 1B), *K152A,* nor *K147A* (Figure 6 – figure supplement 1E) increase the UUG:AUG initiation ratio in otherwise WT cells lacking *SUI5.* These findings suggest that the contacts made by K147 and K152 at the subunit interface are not critically required to stabilize the open, scanning conformation of the PIC and, rather, exclusively stabilize the closed state. The selective effect of *K152E* and *K152A* on the closed complex might involve the fact mentioned above that K152 may make a different pair of rRNA contacts in the open and closed states. For K147, which appears to make the same interactions in both states, it could be proposed that loss of its eIF3c interaction in the open complex conferred by *K147A* is compensated by one or more other interactions in the PIC that uniquely stabilize the open complex.

#### (ii) Evidence that interactions of eIF3b RRM residue Lys-122 and β-propeller residue Lys-346 with rRNA on the 40S interface preferentially stabilize the open conformation of the 48S PIC

eIF3b residue Lys-122 in the RRM appears to contact rRNA helix 24 (Fig. 3-figure Supplement 1A), including residues G985-G986 in the open complex, but only G985 in the closed complex (Table 1). Substituting Lys-122 with Ala, Asp, or Glu, designed to eliminate contact with the rRNA (*K122A*) or replace it with electrostatic repulsion (*K122D* and *K122E*), all conferred dominant lethal phenotypes. This was inferred from our inability to recover transformants containing the corresponding *prt1* mutant alleles even on galactose medium where WT eIF3b/Prt1 is expressed from the chromosomal *P_GAL_-PRT1* allele. Interestingly, whereas Arg substitution of this same residue (*K122R*) has no effect on cell growth, it produces a weak Sui^-^ phenotype, increasing the UUG:AUG *HIS4-lacZ* reporter initiation ratio by ≈1.8-fold (Figure 6C, cols. 1-2). One way to explain these findings is to propose that interaction of K122 with rRNA residues on the 40S interface is essential, such that its replacement with Ala, Asp, or Glu is lethal. The dominance of this lethal phenotype indicates that the mutant protein is expressed and competes with WT eIF3b/Prt1 for incorporation into the eIF3 complex. The basic side-chain introduced by *K122R* would maintain the essential electrostatic attraction with rRNA, but the larger size of the Arg versus Lys side-chain would perturb RRM binding to the rRNA in a manner that diminishes its ability to stabilize the open complex. Consistent with this interpretation, we observed that *K122R* exacerbates the His^+^/Sui^-^ phenotype and elevated *HIS4-lacZ* UUG:AUG ratio conferred by the *sui1-K60E* substitution, which is known to weaken eIF1 binding to the 40S subunit, destabilizing the open complex and increasing inappropriate transition to the closed state at UUG codons (Martin-Marcos et al., 2013) (Figure 6D, rows 2 & 4 vs. 1; and Figure 6E, cols. 2 and 4 vs. 1).

We found that the *K122R* substitution does not confer an Ssu^-^ phenotype in the presence of *SUI5* (Figure 6 – figure supplement 1C), suggesting that it either does not appreciably destabilize the closed complex, or that it confers a relatively greater impairment of the open complex with the net effect of only a weak Sui^-^ phenotype. The relatively greater destabilization of the open complex might be explained by proposing that the *K122R* substitution primarily perturbs K122 possible interaction with G986, found only in the open complex, with minimal effects on the likely G985 interaction common to both conformations.

Interestingly, the Sui^-^ phenotype of *K122R* was found to be suppressed by the *K91E* substitution in the RRM, with the UUG:AUG ratio being returned to the WT level in the *K122R K91E* double mutant (Figure 6C, cols. 2-3). K91 appears to interact with the rRNA residue U1011 in the open complex, and possibly with U1011, A1012, and G986 in the closed complex (Table 1). An intriguing possibility is that *K91E* alters the position of G986 in a manner that overcomes the perturbation of the adjacent rRNA residue G985 by the *K122R* substitution as the means of suppressing the Sui^-^ phenotype of the latter. The *K91E* substitution has no effect on cell growth alone or in combination with *K122R* (Figure 6 – figure supplement 1A, rows 3 & 5 vs. 1), nor does it alter the UUG:AUG ratio in otherwise WT cells (Figure 6 – figure supplement 1E).

Finally, residue K346 in the eIF3b β-propeller approaches rRNA helix 44 (Fig. 3-figure Supplement 1A), possibly contacting residue A1742 on the subunit interface only in the open conformation (Table 1). Consistent with selective destabilization of the open state and concomitant shift to the closed state, we found that K346E confers a weak Sui^-^ phenotype, increasing the UUG:AUG ratio by ≈2.2-fold (Figure 6C, cols. 1 & 4).

#### (iii) Evidence that electrostatic repulsion of eIF3b β-propeller residues Asp397, Glu398, Asp297, and Glu308 with rRNA or eIF2γ on the 40S interface facilitate relocation of the eIF3b-3i-3g-3a-Cter module to the 40S solvent-exposed surface

In addition to observing Ssu^-^ or Sui^-^ phenotypes for substitutions designed to weaken interactions of the eIF3b RRM or β-propeller on the subunit interface, we also examined substitutions that would be expected to stabilize interaction of the β-propeller at the interface, hypothesizing that such substitutions would impede relocation of the eIF3b/eIF3g/eIF3i module back to the solvent side of the 40S subunit. This would be expected to impair start codon recognition, particularly at near-cognate UUG codons, and confer an Ssu^-^ phenotype. Interestingly, the acidic eIF3b residues D397 and E398 are in proximity to helix 44 (Fig. 3-figure Supplement 1A) and may contact the phosphate backbone of rRNA residue G1743 exclusively in the closed complex (Table 1). Mutations that substitute both of these acidic residues with the basic residue Arg confer strong Ssu^-^ phenotypes, completely reversing the elevated UUG:AUG initiation ratio, Slg^-^ at 37°C, and His^+^ phenotypes conferred by *SUI5* (Figure 7A, rows 1 & 3-4; Figure 7B, cols. 1 and 3-4). These findings are consistent with the idea that the adjacent acidic residues D397 and E398 exert electrostatic repulsion with rRNA residue G1743 in the closed complex that facilitates relocation of the WT eIF3b-3i-3g-3a-Cter module back to the 40S solvent side, and that introducing electrostatic attraction between the β-propeller and rRNA at this interface by the D397R and E398R substitution impedes this relocation and subsequent start codon selection.

**Figure 7.**
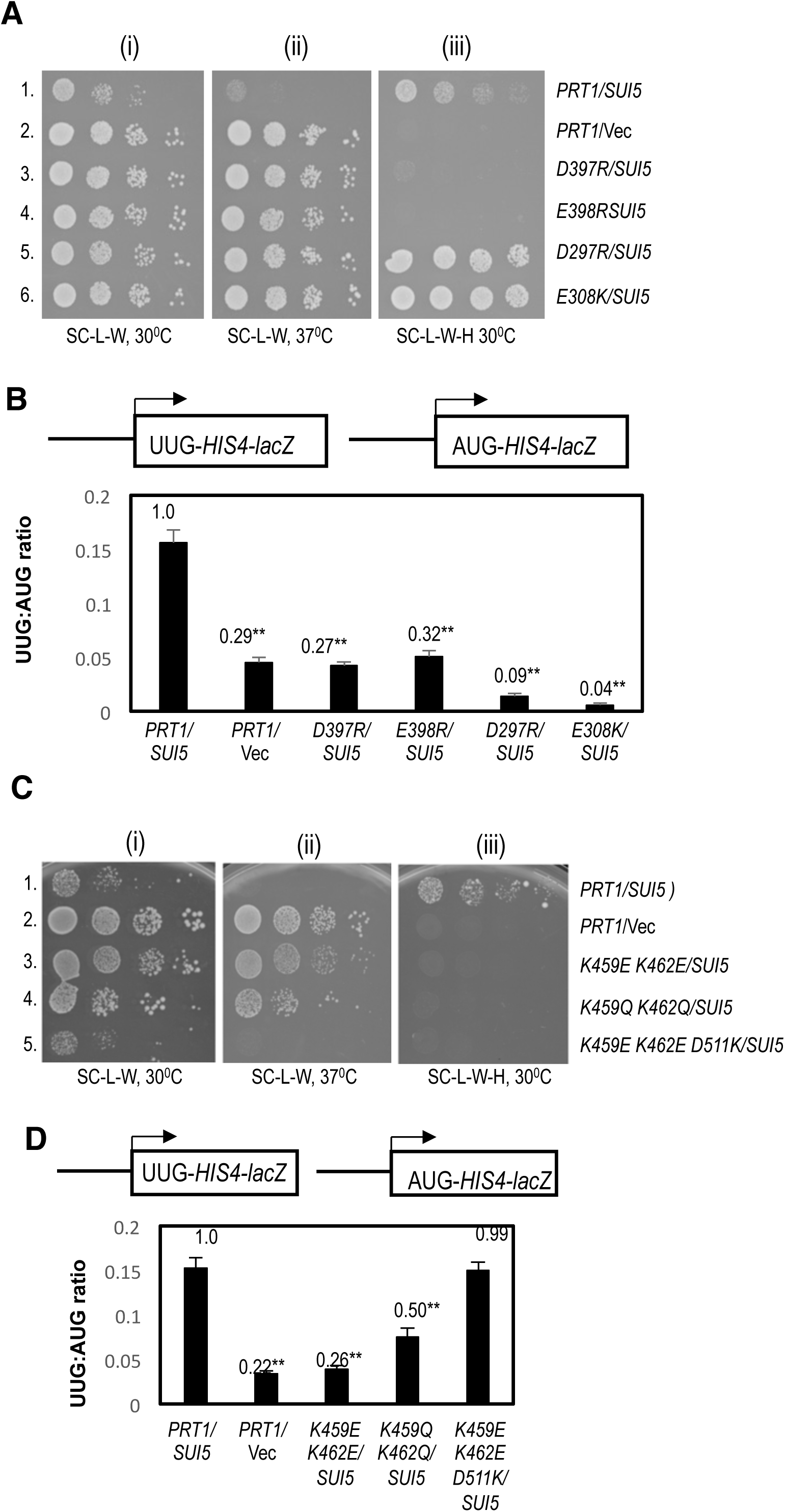
Effect of substitutions in the eIF3b β-propeller on the fidelity of start codon selection in vivo. (A)-(B) Evidence that introducing excessively stable interactions of the eIF3b β-propeller with the 40S subunit or eIF2γ at the subunit interface increases discrimination against UUG start codons. **(A)** Serial dilutions of transformants of *P_GAL1_-PRT1 his4-301* strain HD3607 with the indicated *PRT1* alleles on low-copy (lc) *LEU2* plasmids and either single-copy (sc) *TRP1 SUI5* plasmid p4281 or empty vector YCplac22 (Vec) were spotted on synthetic complete medium lacking Leu and Trp (SC-L-W) or the same medium also lacking His (SC-L-W-H) and incubated at 30°C or 37°C for 3-5d. (B) *HIS4-lacZ* UUG:AUG ratios were determined for strains in (A) exactly as in Figure 6B. **(C)-(D) Evidence that weakening interactions of eIF3b β-propeller residues at the solvent-exposed surface of the 40S subunit increases discrimination against UUG start codons**. **(C)** Serial dilutions of transformants of *P_GAL1_-PRT1 his4-301* strain HD3607 with the indicated *PRT1* alleles on low-copy (lc) *LEU2* plasmids and either single-copy (sc) *TRP1 SUI5* plasmid p4281 or empty vector YCplac22 (Vec) were spotted on synthetic complete medium lacking Leu and Trp (SC-L-W) or the same medium also lacking His (SC-L-W-H) and incubated at 30°C or 37°C for 3-5d. (D) *HIS4-lacZ* UUG:AUG ratios were determined for strains in (C) exactly as in Figure 6B.

Similar findings were made for basic substitutions of two acidic residues in the eIF3b β-propeller, D297 and E308, that are in proximity to acidic residues of eIF2γ in the closed complex, eIF2γ-D460 and eIF2γ-E506, respectively (Fig. 3-figure Supplement 1A & Table 1). Replacing the eIF3b acidic residues with Arg (*D297R*) or Lys (*E308K*) each conferred Ssu^-^ phenotypes in *SUI5* cells, strongly suppressing the UUG:AUG initiation ratio (Figure 7B, cols. rows. 1, 5-6) and Slg^-^ at 37°C (Figure 7A, rows 1, 5-6) conferred by *SUI5*. Unexpectedly, neither mutation reverses the His^+^ phenotype of *SUI5* (Figure 7A, rows 1, 5-6), which might indicate a confounding effect of the mutations on histidine biosynthesis or utilization that we observed previously for certain other suppressors of *SUI5* (Saini et al., 2014). Notwithstanding this last complexity, the findings on *D297R* and *E308K* conform to the model that electrostatic repulsion between the eIF3b β-propeller and eIF2γ serves to facilitate relocation of the WT eIF3b-3i-3g-3a-Cter module to the solvent side of the 40S in a manner that can be impeded by replacing electrostatic repulsion with attraction at the β-propeller:eIF2γ interface. None of the basic substitutions in acidic residues D397, E398, D297, or E308 affected cell growth (Figure 6 – figure supplement 1A, rows 11-14 vs. 9) nor the UUG:AUG ratio in the absence of *SUI5* (Figure 6 – figure supplement 1D).

#### (iv) Evidence that possible interactions of eIF3B β-propeller residues Lys459 and Lys462 with rRNA on the solvent side of the 40S promote the relocation of the eIF3b-3i-3g-3a-Cter module

The genetic results described above provide evidence that eIF3b contacts on the subunit interface are physiologically important for the fidelity of start codon selection. We hypothesized that eIF3b contacts on the solvent side of the 40S would also affect initiation by influencing whether the eIF3b-3i-3g-3a-Cter module can efficiently relocate back to this side of the 40S subunit to allow the completion of start codon selection. Examining the structure of py48S-5N (PDB: 6FYX), in which the eIF3b-3i-3g-3a-Cter module is found on the solvent side of the 40S, revealed that eIF3b β-propeller residues K459 and K462 are in proximity to rRNA residues of h16 on the solvent side of the 40S, with K462 contacting the phosphate backbone of residue U498. In accordance with our hypothesis, the *K459E/K462E* double substitution in the β-propeller confers a strong Ssu^-^ phenotype in *SUI5* cells, suppressing the Slg^-^ at 37°C and His^+^ growth phenotypes and the elevated UUG:AUG ratio conferred by *SUI5* (Figure 7C, rows 1 & 3; Figure 7D, cols. 1 and 3). *K459E/K462E* has no effect on the UUG:AUG initiation ratio in cells lacking *SUI5* (Figure 6 – figure supplement 1E); however, it does confer a moderate Slg^-^ phenotype at 30°C in otherwise WT cells (Figure 6 – figure supplement 1A, row 18), suggesting that this β-propeller:rRNA contact on the solvent side of the 40S might be important for the rate of initiation and not only its fidelity. Reasoning that the Glu substitutions of these eIF3b residues would not only eliminate electrostatic attraction but also introduce electrostatic repulsion with rRNA, we asked whether Gln rather than Glu substitutions have a less severe Ssu^-^ phenotype. Indeed, unlike *K459E/K462E*, the *K459Q/K462Q* substitution conferred no Slg^-^ in otherwise WT cells (Figure 6 – figure supplement 1A, row 20 vs. 18 & 15) and suppression of the elevated UUG:AUG ratio conferred by *SUI5* was less complete for *K459Q/K462Q* compared to the *K459E/K462E* mutant (Figure 7D, rows 1 and 4 vs. 3); although still strong enough to suppress the His^+^ and Slg-phenotypes of *SUI5* (Figure 7C, rows 1 & 4). These findings support the idea that electrostatic attraction between eIF3b residues *K459/K462* and rRNA residues on the solvent side of the 40S promote relocation of the eIF3b-3i-3g-3a-Cter module back to the solvent side to complete the process of start codon selection.

Interestingly, when *K459E/K462E* are combined with *D511K*, the ability of K459E/K462E to suppress the Slg^-^ at 37°C and elevated UUG:AUG initiation ratio conferred by *SUI5* is lost, as both the Slg^-^ at 37°C and elevated UUG initiation conferred by *SUI5* are reinstated in the triple mutant (Figure 7C, row 1 & 3 vs. 5; Figure 7D, col. 1 & 3 vs. 6). While it appears that the His^+^ phenotype of *SUI5* was not reinstated, the absence of a strong His^+^ phenotype in the *K459E/K462E/D511K* triple mutant in Figure 7C (row 5) can be explained by the strong Slg^-^ phenotypes conferred on +His medium by this triple mutation compared to the *K459E/K462E* double mutation (row 5 vs. 3). While not in direct contact, D511 is in proximity to 18S rRNA residue C499 of h16 (PDB: 6FYX), such that introducing lysine at this position might strengthen binding of the β-propeller to the solvent surface of the 40S and counteract the predicted effects of *K459E/K462E* in weakening this same interaction. If so, this would restore both efficient relocation of the eIF3b-3i-3g-3a-Cter module back to the solvent side of the 40S and efficient start codon selection, and thereby reverse the effects of *K459E/K462E* in reducing UUG initiation.

## Concluding remarks

The structures of py48S-open-eIF3 PIC and py48S-closed-eIF3 presented here show the interactions of the β-propeller and RRM of eIF3b at the 40S subunit interface. eIF3b adopts a similar conformation in both complexes with subtle differences in its interactions with 48S PIC components at the subunit interface in the open versus closed states (Figure 5C, Movie 1). eIF3i is also resolved (especially in the py48S-closed-eIF3 map) in the new structures; however, its β-propeller is weakly attached to eIF3b and does not make any direct contacts with the 40S subunit. This stands in contrast to a model proposed recently for the rearrangement of eIF3 during translation initiation wherein the β-propellers of eIF3b and eIF3i re-locate independently to the subunit interface at different stages of the process (Simonetti et al., 2016). This model was based on a model of a mammalian 48S late stage IC in which the β-propeller of eIF3i was positioned at the GTPase-binding site on the subunit interface (Simonetti et al., 2016). However, as this density was later re-interpreted as the ABCE1 protein (Mancera-Martínez et al., 2017), the model proposed by Simonetti et al. (Simonetti et al., 2016) now seems unlikely. Instead, we propose an alternative model in which the β-propellers of eIF3b and eIF3i re-locate together from the solvent interface of the 43S PIC to the subunit interface after mRNA binding or in early stages of codon:anticodon recognition, with only the eIF3b propeller contacting the 40S subunit in both states, and thereafter these domains relocate together back to their original positions on the solvent interface after the release of eIF1 and its replacement by the eIF5 N-terminal domain at the subunit interface (Llácer et al., 2018) (Figure 8A,B).

**Figure 8.**
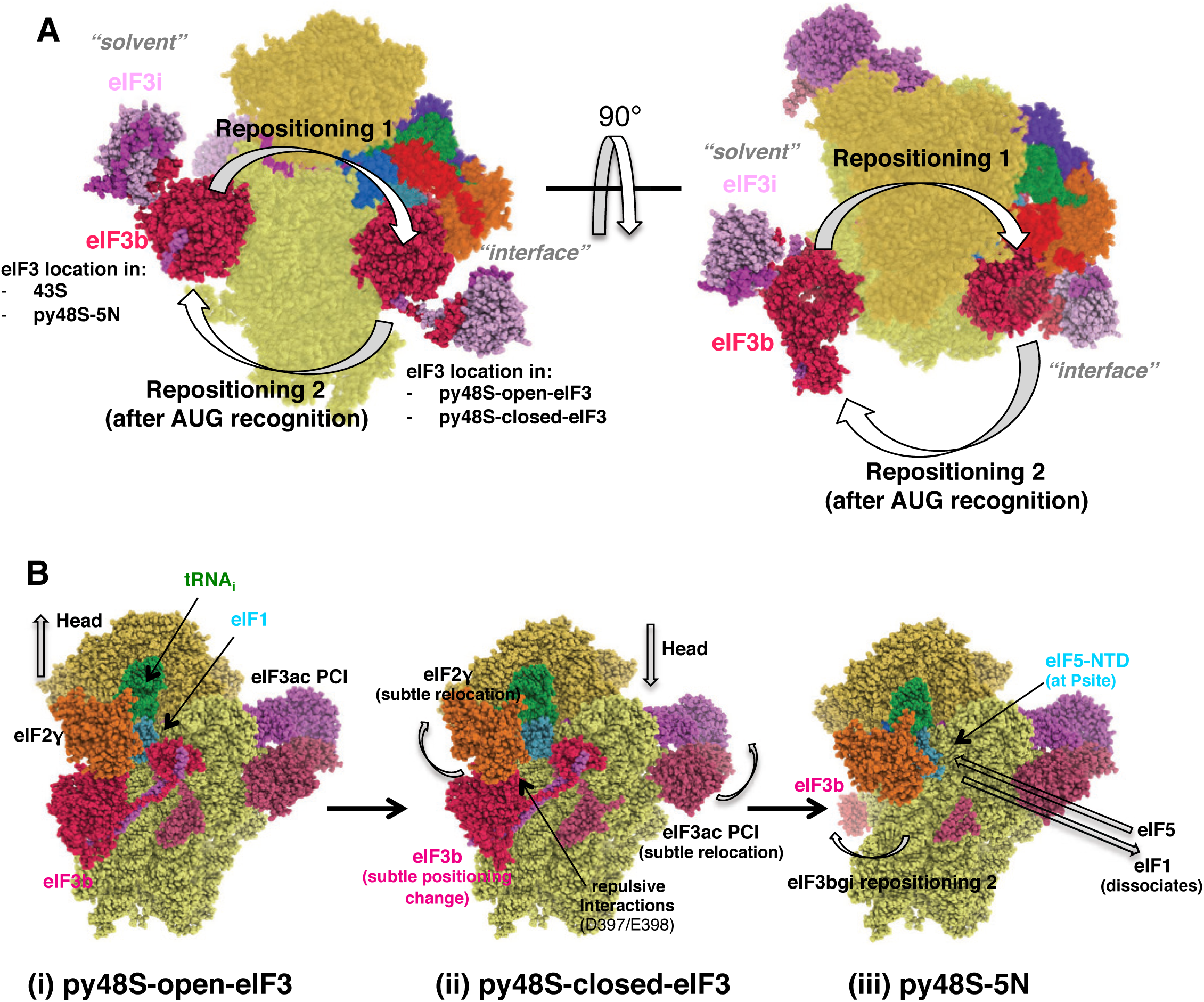
Model depicting reversible repositioning of the eIF3b/eIF3i/eIF3g/eIF3a-Cterm module between the solvent-exposed and subunit-interface surfaces of the 40S at the onset of scanning and following AUG recognition. **(A)** Side and top views of a composite representation showing the two different locations observed for the eIF3b/eIF3i/eIF3g/eIF3a-Cterm module on the solvent or subunit surfaces of the 40S subunit in 43S/py48S-eIF5N and py48S-open-eIF3/py48S-closed-eIF3, respectively (Llácer et al., 2018). The arrows depict the direction of the movement of this eIF3 subcomplex from the solvent surface to the subunit interface upon 43S PIC attachment to mRNA and formation of the open, scanning conformation of the 48S PIC (Repositioning 1), and then back to the solvent surface after AUG recognition, complete accommodation of tRNA_i_ and eIF1 dissociation from the 48S complex (Repositioning 2). **(B)** Representation showing the subtle conformational changes of different eIF3 elements, tRNA, the 40S head and eIF2γ at the subunit interface accompanying transition from the open to closed states of the 48S complex on AUG recognition (i)-(ii); followed by repositioning of the eIF3b/eIF3i/eIF3g/eIF3a-Cterm module (eIF3bgi) to the solvent side of the 40S following AUG recognition, eIF1 dissociation and replacement of eIF1 with the eIF5 NTD near the P-site (iii). For clarity, eIF3i/g, eIF3a-CTD eIF3c-NTD, and eIF2αβ are not shown.

We found that genetic perturbation of interactions of eIF3b with the 48S subunit interface reduce the fidelity of start codon recognition in vivo, indicating that eIF3b contacts on the subunit interface are physiologically important for stringent selection of AUG start codons. Certain substitutions affecting eIF3b interactions at the subunit interface modestly increase the probability of selecting near-cognate UUG codons (Sui^-^ phenotype), whereas others have the opposite effect on fidelity in strongly reducing initiation at UUG codons (Ssu^-^ phenotype). These alternative outcomes have been observed previously for mutations in eIF1 and eIF1A that specifically disfavor the open (Sui^-^) or closed (Ssu^-^) conformation of the 48S PIC, indicating that preferentially perturbing the open complex allows inappropriate rearrangement from the scanning conformation to the closed state at UUG codons (Sui^-^ phenotype), whereas preferentially destabilizing the closed state shifts the system to the open conformation to impair selection of UUG codons (Ssu^-^ phenotype) (Hinnebusch, 2017). The mismatched duplex formed by Met-tRNA_i_ at UUG codons appears to render them more sensitive than AUG codons to perturbations of the closed state and thereby lowers the UUG:AUG initiation ratio in Ssu^-^ mutants. The modest Sui^-^ phenotype conferred by two substitutions in the RRM (K122R) and β-propeller (K346E) (Figure 3 figure Supplement 1A) suggest that the predicted interactions of these residues with rRNA at the subunit interface preferentially stabilize the open, scanning conformation of the PIC. The much stronger Ssu^-^ phenotypes conferred by the β-propeller substitutions K147E and K152E identifies an especially critical role for these rRNA contacts in achieving or stabilizing the closed conformation of the PIC required for start codon recognition (Fig. 3-figure Supplement 1A). The fact that perturbing different eIF3b contacts at the subunit interface can have opposing effects on fidelity can be explained by proposing that the affected interactions are either restricted to one conformation, owing to the subtle changes in position of the eIF3b-3i-3g-3a-Cter module between the open and closed conformations of the scanning PIC (Movie 1 and Fig. 8C(i)-(ii)), or they are relatively more important in stabilizing one of the two conformations.

Interestingly, equally strong Ssu^-^ phenotypes were observed for eIF3b β-propeller substitutions expected to impede relocation of the eIF3b-3i-3g-3a-Cter module from the interface back to the solvent side of the 40S subunit, by either of two mechanisms. The first involves β-propeller substitutions designed to diminish electrostatic repulsion between eIF3b and rRNA (D397R and E398R) or eIF3b and eIF2γ (D297R and E308K) occurring at the subunit interface of wild-type PICs, which should engender an abnormally stable interaction of the eIF3b-3i-3g-3a-Cter module at the subunit interface (Figure 3 figure Supplement 1A). The second mechanism involves a double substitution predicted to disrupt eIF3b interaction with rRNA specifically on the solvent side of the 40S, K459E/K462E, which should selectively weaken association of the eIF3b-3i-3g-3a-Cter module on the solvent surface (Figure 3 figure Supplement 1A). The repulsive interactions between eIF3b residues D397/E398 and rRNA at the subunit interface of wild-type PICs, which should favour release of the eIF3b-3i-3g-3a-Cter module for its relocation back to the solvent surface on AUG recognition (perturbed by the first mechanism), appear to exist only in the closed conformation of the PIC, owing to downward movement of the 40S head in the transition from open to closed states (Figure 8B (ii) vs. (i)).

Together, our genetic findings suggest that efficient start codon recognition, and utilization of near-cognate UUG codons in particular, depends not only on interactions of eIF3b at the 40S subunit interface that preferentially stabilize the closed conformation of the PIC, but also on the timely dissolution of these interactions to permit relocation of the eIF3b-3i-3g-3a-Cter module back to the solvent side of the 40S subunit following AUG recognition. As such, different eIF3b substitutions predicted to either selectively weaken its interactions on the subunit interface, to render these interactions excessively stable, or to selectively diminish its interactions on the solvent side of the 40S subunit, all have the same consequence of strongly disfavouring initiation at near-cognate UUG codons. Other eIF3b substitutions that appear to preferentially destabilize its subunit-interface interactions in the open conformation of the PIC confer the opposite effect of enhancing UUG initiation; but judging by the relative weakness of these latter phenotypes, those perturbations have a decidedly smaller impact on fidelity compared to those that preferentially destabilize the closed conformation of the scanning PIC.

The relocation of the eIF3b-3i-3g-3a-Cter module from the 40S subunit interface back to the solvent side of the 40S (Repositioning 2, Figure 8A) may facilitate dissociation of eIF1 from the 40S interface owing to loss of eIF1 interactions with the eIF3b RRM, or promote subsequent dissociation of eIF2 from Met-tRNA_i_ owing to loss of eIF2γ contacts with the eIF3b β-propeller occurring at the subunit interface. This relocation should also be required for joining of the 60S subunit, which requires extensive interactions with the 40S interface, some of which would be sterically hindered by the presence of the eIF3b-3i-3g-3a-Cter module at this position.

eIF3 is involved in nearly all steps of initiation, including stimulating the binding of eIF2 TC and other eIFs to the 40S ribosomal subunit; the attachment of the 43S complex to mRNA; the subsequent scanning along mRNA to reach the start AUG codon; and finally, prevention of the joining of small and large ribosomal subunits prior to start codon recognition (Hinnebusch, 2014; Hinnebusch, 2017; Valášek et al., 2017). However, little is known about the molecular basis of these various functions of eIF3 and the role of each subunit in executing them. In particular, there was no understanding of the functional significance of the conformation of eIF3 in the PIC with its peripheral eIF3b-3i-3g-3a-Cter module positioned on the subunit interface, as described in detail here. Overall, the structural models of eIF3 and genetic analysis presented in this study suggest that distinct eIF3b interactions on the subunit interface play crucial roles in stabilizing one or more features of the closed conformation, or in modulating relocation of the eIF3b-3g-3i module back to the solvent side of the 40S to license one or more late steps of the initiation pathway following recognition of the start codon.

## MATERIALS & METHODS

### Reconstitution of the 48S complex

*Kluyveromyces lactis* 40S subunits were prepared as described earlier (Fernández et al., 2014). *Saccharomyces cerevisiae* eIF2 was expressed in yeast while eIF1, eIF1A, eIF5, eIF4A, eIF4B and the eIF4G1:eIF4E complex were expressed in *Escherichia coli* as recombinant proteins and purified as described (Acker et al., 2007) (Mitchell et al., 2010) (Llácer et al., 2018). eIF3 was also expressed in yeast as previously described (Acker et al., 2007) but with modifications to avoid eIF5 contamination. To achieve the latter, we replaced the phosphocellulose column with a Q-sepharose column and eluted it with an extended KCl gradient from 100 mM to 1M (50 column volumes), obtaining two batches of eIF3, one without eIF5, the other enriched in eIF5. Wild type tRNA_i_ was overexpressed and purified from yeast and aminoacylated as described (Acker et al., 2007). The mRNA expression construct comprised a T7 promoter followed by the 49-nt unstructured mRNA sequence of 5′-GGG[CU]_3_[UC]_4_UAACUAUAAAA<u>AUC</u>[UC]_2_UUC[UC]_4_GAU-3′ (with start codon underlined), cloned between XhoI and NcoI sites in a pEX-A2 plasmid (Eurofins Genomics). mRNA was purified and capped as in (Llácer et al., 2018).

The 43S PIC was reconstituted first, by incubating 40S with eIF1, eIF1A, TC (consisting of eIF2, GDPCP and Met-tRNA_i_), and eIF3 in 40S:eIF1:eIF1A:TC:eIF3 molar ratios of 1:2.5:2.5:2:2, in 20 mM MES, pH 6.5, 80 mM potassium acetate, 10 mM ammonium acetate, 5-8mM magnesium acetate, 2mM dithiothreitol (DTT), 1 µM zinc acetate. Separately, an mRNA-eIF4 complex was prepared, containing eIF4G1, eIF4E, eIF4A, eIF4B and capped mRNA in molar ratios of 1.5:1.5:5:2:2.5 with respect to the 40S ribosome, in 20 mM Hepes, pH 7.4, 100 mM potassium chloride, 5mM magnesium chloride, 2mM DTT, 3mM AMPPNP). The volume of the mRNA-eIF4 reaction mixture was 5 times smaller than that for the 43S PIC. Both the 43S mixture and the mRNA-eIF4 mixture were incubated separately for 5 min at room temperature before combining them. After incubation for 2 min at room temperature, the sample (at a 40S concentration of 80 nM) was cooled to 4°C and used immediately to make cryo-EM grids without further purification.

### Electron microscopy

Three µl of the 48S complex was applied to glow-discharged Quantifoil R2/2 cryo-EM grids covered with continuous carbon (of ∼50Å thick) at 4 °C and 100% ambient humidity. After 30 s incubation, the grids were blotted for 2.5-3 s and vitrified in liquid ethane using a Vitrobot Mk3 (FEI). Automated data acquisition was done using the EPU software (FEI) on a Titan Krios microscope operated at 300 kV under low-dose conditions (30 e-/Å2) using a defocus range of 1.2 – 3.2 µm. Images of 1.1 s/exposure and 34 movie frames were recorded on a Falcon III direct electron detector (FEI) at a calibrated magnification of 104,478 (yielding a pixel size of 1.34 Å). Micrographs that showed noticeable signs of astigmatism or drift were discarded.

### Analysis and structure determination

The movie frames were aligned with MOTIONCORR (Li et al., 2013) for whole-image motion correction. Contrast transfer function parameters for the micrographs were estimated using Gctf (Zhang, 2016). Particles were picked using RELION (Scheres, 2012). References for template-based particle picking (Scheres, 2015) were obtained from 2D class averages that were calculated from particles picked with EMAN2 (Tang et al., 2007) from a subset of the micrographs. 2D class averaging, 3D classification and refinements were done using RELION-2 (Scheres, 2012). Both movie processing (Bai et al., 2013) in RELION-2 and particle ‘‘polishing’’ was performed for all selected particles for 3D refinement. Resolutions reported here are based on the gold-standard FSC = 0.143 criterion (Scheres and Chen, 2012). All maps were further processed for the modulation transfer function of the detector, and sharpened (Rosenthal and Henderson, 2003). Local resolution was estimated using Relion and ResMap (Kucukelbir et al., 2014).

For the py48S-open-eIF3 dataset, 2610 images were recorded from two independent data acquisition sessions. An initial reconstruction was made from all selected particles (360,729) after 2D class averaging using the yeast 40S crystal structure (PDB: 4V88), low-pass filtered to 60 Å, as an initial model. Next, a 3D classification into 12 classes with fine angular sampling and local searches was performed to remove abnormal or empty 40S particles from the data. Three highly populated classes (144,292 particles) showed density for the TC. Two consecutive masked 3D classifications with subtraction of the residual signal (Bai et al., 2015) were then performed, by creating masks around the density attributed to the TC and to the different domains of eIF3 at the subunit interface. In the first ‘focused’ 3D classification using the TC mask we isolated 104,792 particles containing a distinct density for the TC. In the second round of ‘focused’ 3D classification using the eIF3 mask, only 1 class contained eIF3 in high occupancy (13,038 particles, 2.6 % of the total, 4.6Å). The latter class was further divided into two different classes: a) A 48S PIC in a partially open form (7,288 particles; 5.6Å), where the head of the 40S adopts an intermediate position between that in the closed and open forms; b) A 48S PIC in the fully open form (5,750 particles, 5.15Å), which is conformationally identical to our previously reported py48S-open complex. The quality of the density for eIF3 at the subunit interface is better in the latter class than that in the partially open form or one that resulted from the combination of the two classes.

We have also reprocessed our previous dataset of the py48S-closed complex using image processing tools that were not available at the time we published our previous work with the aim of improving the local resolution of eIF3 at the subunit interface by isolating a larger number of particles containing higher occupancy for eIF3. We started with more than 1,182,000 particles after 2D-classification and after an initial reconstruction done with all selected particles, a first attempt of focused classification with subtraction of the residual signal using a mask around the observed eIF3 beta propeller near the TC did not yield a satisfactory result. Instead, we carried out a conventional 3D-classification in 16 classes and selected the classes containing a clear density for the TC (365,343) followed by movie-processing/particle polishing. Then we performed three consecutive masked 3D classifications with subtraction of the residual signal, by using masks around the density attributed to the eIF3 beta propeller near the TC (‘bgi mask’; selecting 1 class out of four, 128,915 particles), around the tRNA/alpha subunit of eIF2 (‘tRNA/alpha mask’; selecting 3 classes out of five, 55,915 particles) and around all the different domains of eIF3 at the subunit interface (‘eIF3 mask’). In this latter classification we obtained three different classes in which only one of them (12,586 particles, 5.75Å) showed a very distinct an unambiguous density for all the eIF3 subunits, reflecting the higher mobility and flexibility of all these eIF3 domains.

### Model building and refinement

The previous atomic model of py48S in open conformation (PDB: 3JAQ) was placed into density by rigid-body fitting using Chimera (Pettersen et al., 2004). Overall densities for eIF3 at the subunit interface were similar in both the open and closed structures, so model building for eIF3 was done simultaneously in both maps, using one or another depending on the quality of the map for each of the different domains of eIF3. Initially, all the different domains of eIF3 were fitted by rigid-body using Chimera as follows:

a. The eIF3a/eIF3c PCI dimer was taken from the PDB:3JAP and placed in its corresponding density at the solvent side of the 40S in the py48S-open-eIF3 map.
b. The eIF3b β-propeller was taken from the PDB:4NOX and placed into the drum-like density below the TC, in almost the same orientation to that found in the PDB:5K1H.
c. The ternary complex eIF3i/eIF3g/eIF3b-cterm originally placed into the above mentioned drum-like density was now placed (together with the helical linker connecting the eIF3b β-propeller with its c-terminal helix in contact with eIF3i) into the low-resolution density in close contact with the eIF3b β-propeller, thereby placing the two beta propellers in a rather similar arrangement to that found at the solvent side of the 40S (Llácer et al., 2018). For a better fitting of eIF3i, this low-resolution area of the map was de-noised using LAFTER (Ramlaul et al., 2019).
d. The eIF3b RRM and the linker connecting it with the eIF3b β-propeller were taken from PDB:5K1H, and placed almost identically, with minor changes (see below).
e. The eIF3a c-term helix in contact with eIF3b and the eIF3c helical bundle were not displaced from their original positions.

Then, each domain of eIF3 was rigid-body fitted in Coot (Emsley et al., 2010) and further model building was also done in Coot v0.8, which included replacement of each amino acid of eIF3b by its counterpart in *S. cerevisiae*; and eIF3a C-term helix model building. Possible residue numbering for the eIF3a C-term helix was based on eIF3a secondary structure predictions (consensus of programs PSSPred, SOPMA, Jpred) and sequence conservation among homologs from an alignment done in Clustal Omega using *S. cerevisiae, H. sapiens, A. thaliana, C. elegans, D. melanogaster,* and *S. pombe* sequences (with the expectation of higher conservation in regions of eIF3a involved in interactions), distance restraints between this helix and the PCI domain of eIF3a at the solvent side (also taking into account sequence conservation in different organisms) and known interactions with eIF3b and the eIF3b RRM domain (Chiu et al., 2010) and with the ribosome (Szamecz et al., 2008).

### eIF3c N-term model building

For the 5-helical bundle we have used blurred maps of our previous py48S-closed model. Length of the helices matched the lengths predicted by secondary structure prediction programs and the newer improved maps helped to model the connection between helices, aided by the fact that the extra density connecting the helical-bundle with eIF1/RRM and the PCI helped identifying the N- and C-terminus, respectively, of the helical-bundle. Density for some bulky side chains also helped in the sequence assignment as well as sequence conservation among homologs from an alignment of *S. cerevisiae, H. sapiens, A. thaliana, D rerio, D. melanogaster,* and *S. pombe* sequences (again with the expectation that residues of eIF3c involved in interactions are more highly conserved).

Model refinement was carried out in Refmac v5.8 optimized for electron microscopy (Brown et al., 2015), using external restraints generated by ProSMART and LIBG (Brown et al., 2015). Average FSC was monitored during refinement. The final model was validated using MolProbity (Chen et al., 2010). Cross-validation against overfitting was calculated as previously described (Brown et al., 2015) (Amunts et al., 2014). Buried surface areas and contacts between eIF3b and other 48S components were calculated using PISA (Krissinel and Henrick, 2007). All figures were generated using PyMOL (DeLano, 2006) Coot or Chimera.

### Plasmid constructions

Plasmids employed in this work are listed in Table S3. Plasmids pDH14-29, pDH14-90, pDH15-26, pDH15-67, pDH14-95, pDH14-96, pDH14-56, pDH14-57, pDH15-9, pDH15-10, pDH14-61, pDH15-11, pDH15-76, and pDH15-42 were derived from the low copy number (lc) *LEU2 PRT1* plasmid p5188 by site-directed mutagenesis of *PRT1* using the QuickChange XL kit (Agilent Technologies) and the corresponding primers in Table S2. DNA sequence changes were verified by sequencing the entire *PRT1* gene. Plasmids pDH15-59 and pDH15-61 were derived from p5188 and pDH14-90, respectively, by employing the marker swap plasmid pLT11 as described (Cross, 1997), and verified by sequencing.

### Yeast strain construction

Yeast strains employed in this work are listed in Table S4. Strain HD3607 was derived from H2995 by replacing the promoter of chromosomal *PRT1* with the *GAL1* promoter by one-step gene replacement, as follows: Primers (i) 5’GAA AAT GCT CCA GTG GCT ACG AAT GCA AAC GCT ACC ACT GAC CAA GAG GGT GAT ATT CAC CTA GAA TAG GAA TTC GAG CTC GTT TAA AC and (ii) 5’ CTA TGT ACT GAG ATG TTA TTG AAA TAT AAT TCG TAA ATA TTT TTC AAT GTG CGT GGA AGA AAA TTT T CAT TTT GAG ATC CGG GTT TT were used to amplify by PCR the appropriate DNA fragment from plasmid pFA6a-*kanMX6-P_GAL1_* (p3218), which was used to transform strain HD2955 to Kan^R^, resulting in HD3607. The presence of *P_GAL1_-PRT1* in HD3607 was verified by demonstrating that the lethality on glucose medium can be complemented by lc *PRT1 LEU2* plasmid p5188 but not by an empty vector, and further verified by colony PCR analysis (online protocol, Thermo Fisher Scientific) of chromosomal DNA with the appropriate primers.

Strains HD3648, HD3687, HD3649, HD3848, HD4059, HD4262, HD4256, HD3850, HD3668, HD3669, HD3861, HD3862, HD3672, HD3955, HD4263, and HD4075 were derived from HD3607 by introducing a *LEU2* plasmid containing the indicated mutated *prt1* allele, or empty vector.

Strains HD4081, HD4082, HD4269. HD3995, HD4257, HD4280, HD4281, HD3749, HD4279, HD4012, HD4268, HD4009, and HD3993 were derived from above *PRT1* or *prt1* mutant strains by introducing the sc *TRP1 SUI5* plasmid p4281, or empty *TRP1* vector YCplac22, as indicated in Table S4.

Strain HD4053 was derived from H3956 by replacing the promoter of chromosomal *PRT1* with the *GAL1* promoter by one-step gene replacement, and verified by complementation testing and colony PCR, as described above for strain HD3607. Strains HD4108 and HD4109 were derived from HD4053 by plasmid shuffling to replace the resident lc *URA3 SUI1* plasmid with a lc *LEU2* plasmid containing the *SUI1 or sui1-*K60E alleles. HD4192, HD4193, HD4196, and HD4197 were derived from HD4108 or HD4109 by introducing lc *TRP1* plasmids carrying *PRT1* (pDH15-59) or *prt1-K122R* (pDH15-61), respectively.

### *β*-galactosidase assays

Assays of *β*-galactosidase activity in whole-cell extracts (WCEs) were performed as described previously(Moehle and Hinnebusch, 1991).

### Data Resources

Two maps have been deposited in the EMDB with accession codes EMDB: 0057 and EMDB: 0058, for the py48S-open-eIF3 and py48S-closed-eIF3 maps, respectively. Two atomic coordinate models have been deposited in the PDB with accession codes PDB: 6GSM and PDB: 6GSN, for the py48S-open-eIF3 and py48S-closed-eIF3 models, respectively. These models replace previous PDB:3JAQ and PDB:3JAP models (Llácer et al., 2015) and are also linked to previous published maps EMDB: 3050 and EMDB: 3049, respectively (Llácer et al., 2015).

## Supplemental information

Supplemental information includes four tables and five figures.

## ACKNOWLEDGEMENTS

We are thankful to V. Ramakrishnan at MRC Laboratory of Molecular Biology for his continuous support. The initial phases of this work was carried out in his laboratory. We are also grateful for the technical support with cryo-EM data collection at MRC Laboratory of Molecular Biology. This work was funded by grants from the Spanish government (BFU2017-85814-P), Generalitat Valenciana (SEJI/2019/030) to JLL; by start-up funds [R(IV)090/1076/2017-4252] from Indian Institute of Science, Bangalore, India and the WellcomeTrust/DBT India Alliance Fellowship [IA/I/17/2/503313] to TH; the Human Frontiers in Science Program (RGP-0028/2009) and by the Intramural Research Program of the NIH to AGH. TH also thanks DST-FIST [SR/FST/LS11-036/2014(C)], UGC-SAP [F.4.13/2018/DRS-III (SAP-II)] and DBT-IISc Partnership Program Phase-II (BT/PR27952-INF/22/212/2018) for infrastructure and financial support.

## Supplementary Figures

**Figure 1 – figure supplement 1.**
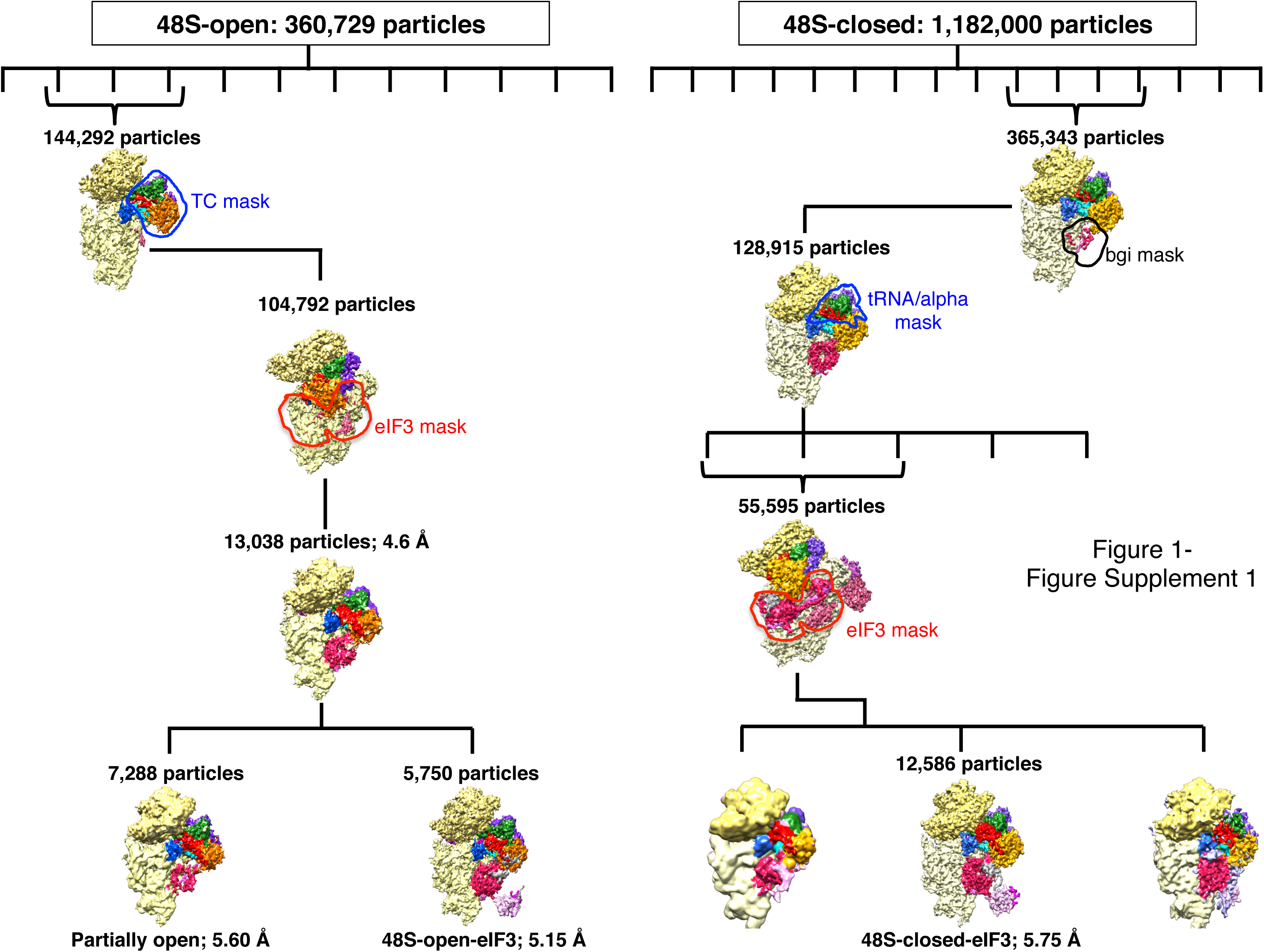
Scheme of 3D classification of data. For py48S-open-eIF3 (*left*), 360,729 particles were selected after 2D classification and an initial 3D refinement was done. After a 3D-classification into twelve different classes, three classes contained clear density for TC and were grouped together (144,292 particles). Then focused 3D classifications were carried out, using separately a ‘TC mask’ first (selecting 104,792 particles) and then an ‘eIF3 mask’ (shown by outlines). As a result, we obtained a class with high occupancy for eIF3 at the subunit interface (13,038 particles) that could be further divided into two classes showing a different degree of head tilting. One of them, dubbed py48S-open-eIF3, corresponds to the fully open head conformation (5,750 particles, at 5.15 Å overall resolution). (*right*) The previous py48S-closed dataset was reprocessed, as follows. 1,182,041 particles were selected after 2D classification and an initial 3D refinement was done. After a 3D-classification into sixteen classes, four classes contained clear density for TC and were joined together. Focused 3D classifications were then carried out. The eIF3 masks ‘bgi mask’ and ‘eIF3 mask’ as well as the ‘tRNA/alpha mask’ used successively for focused 3D classifications are shown in outline. A final class of 12,586 particles showing higher occupancy for eIF3 was obtained, dubbed py48S-closed-eIF3 at 5.75 Å overall resolution. See ‘Analysis and structure determination’ section of Methods for additional details.

**Figure 1 – figure supplement 2.**
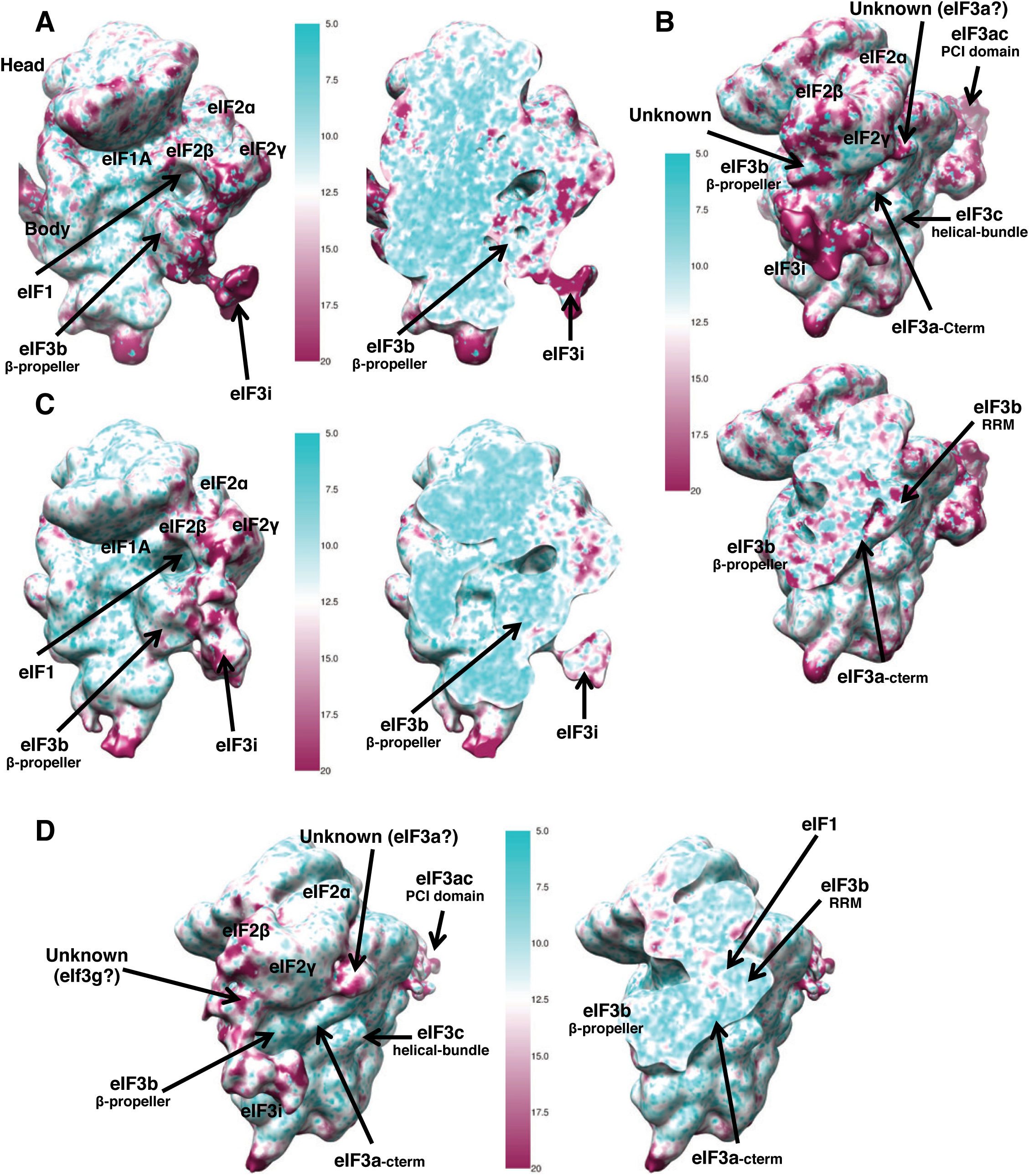
Map quality and local resolution. Surface (left or top) and cross-sections (right or bottom) of gaussian-filtered maps, colored according to local resolution. (A) py48S-open-eIF3, front view (B) py48S-open-eIF3, lateral view (C) py48S-closed-eIF3, front view (D) py48S-closed-eIF3, lateral view

**Figure 1 – figure supplement 3.**
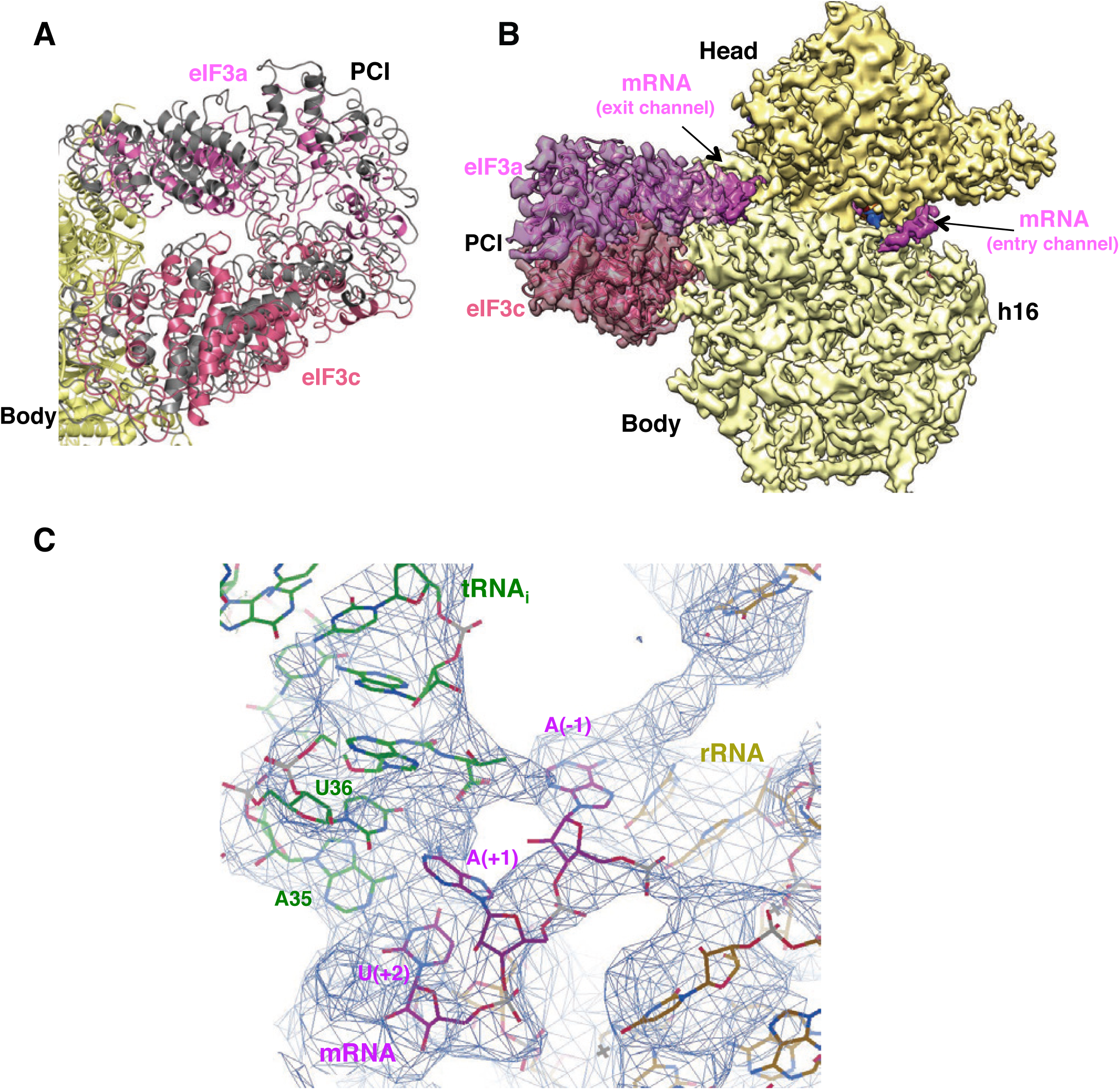
eIF3 PCI conformations and mRNA path in py48S-open-eIF3. (A) Superimposition of the PCI domains in the open/closed maps. (B) py48S-open-eIF3 map, showing density for mRNA at both the entry and exit portions of the mRNA-binding channel on the 40S (C) Detailed view of the codon-anticodon helix of py48S-open-eIF3. There is only density for the first two bases of the codon, A(+1) and U(+2), and the preceding base A(-1).

**Figure 3 – figure supplement 1.**
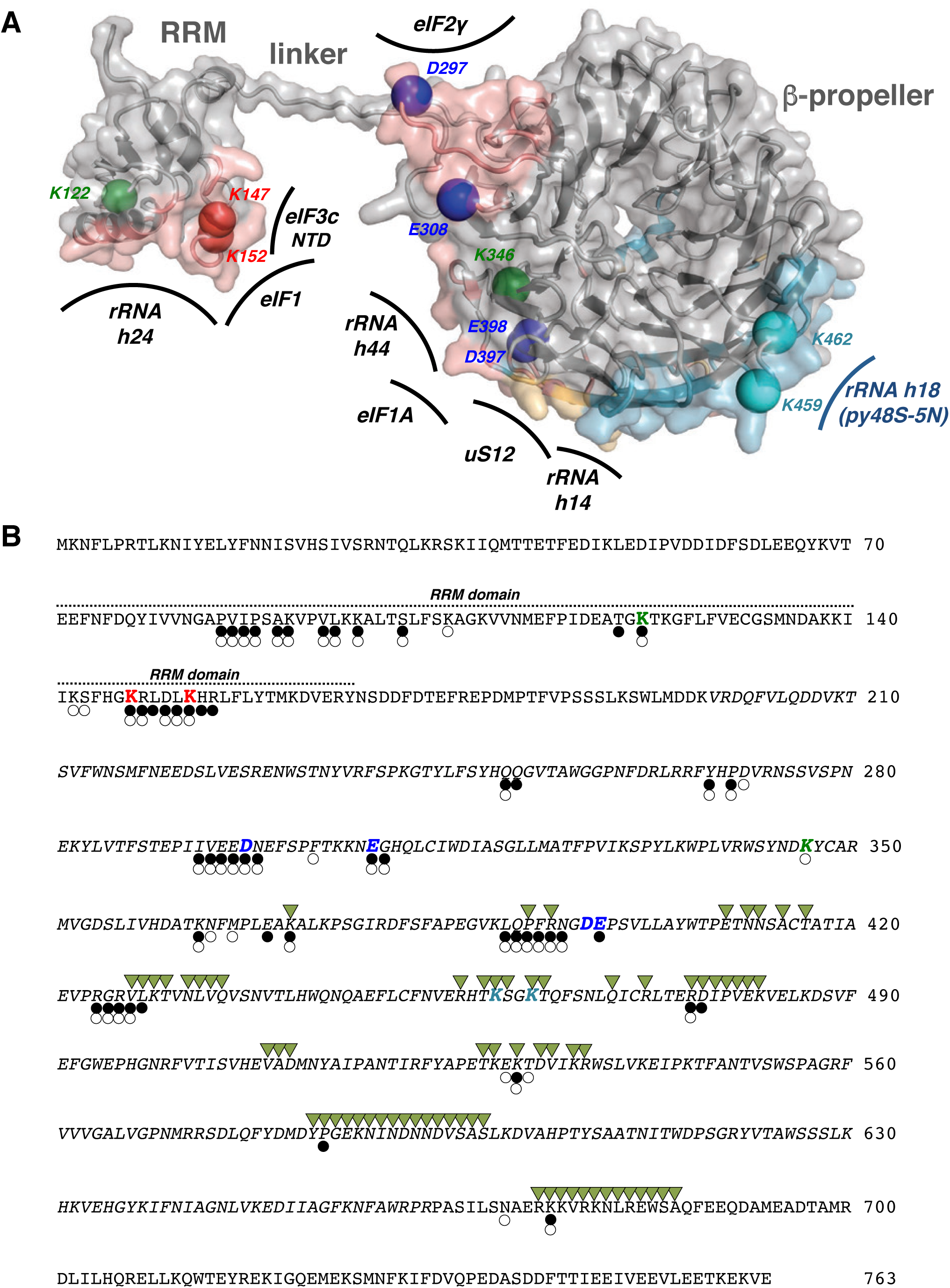
eIF3b contacts and location on the structure of the substituted residues used on the genetics study. (A) Cartoon and transparent surface representation of eIF3b in py48S-open-eIF3. Surfaces colored in salmon encompass residues interacting with the 40S, eIF1, eIF3c or eIF2γ in either py48S-open-eIF3 or py48S-closed-eIF3; surface colored in cyan encompasses residues interacting with the 40S in py48S-5N (PDB: 6FYX); surface colored in yellow includes residues involved in interactions at both the subunit interface and solvent side of the 40S in either py48S-open-eIF3 or py48S-closed-eIF3 and also py48S-5N. Residues substituted in genetic studies are shown as spheres. Based on the phenotypes of the substitutions, red spheres correspond to RRM residues that preferentially stabilize the closed conformation of the 48S PIC, green spheres correspond to residues preferentially stabilizing the open conformation of the 48S PIC, blue and cyan spheres correspond to residues facilitating relocation of the eIF3b/eIF3g/eIF3i module to the 40S solvent-exposed surface either through repulsive interactions with rRNA or eIF2γ at the subunit interface (blue) or attractive interactions with rRNA on the solvent side of the 40S (cyan). Different parts of eIF3b, as well as its interaction partners at the 40S-subunit-interface are labeled. (B) Amino acid sequence of yeast eIF3b/Prt1. Residues substituted in genetic studies are in bold and colored as in (A). Residues in italics belong to the β-propeller of eIF3b. Residues having decreased accessibility [analyzed using a water probe of 1.4A in PISA (Krissinel and Henrick, 2007)] upon binding to 40S/eIF1/eIF3c/eIF2γ in py48S-closed-eIF3 (black circles), py48S-open-eIF3 (transparent circles), or to 40S in py48S-5N (PDB: 6FYX; green triangles) are indicated.

**Figure 6 – figure supplement 1.**
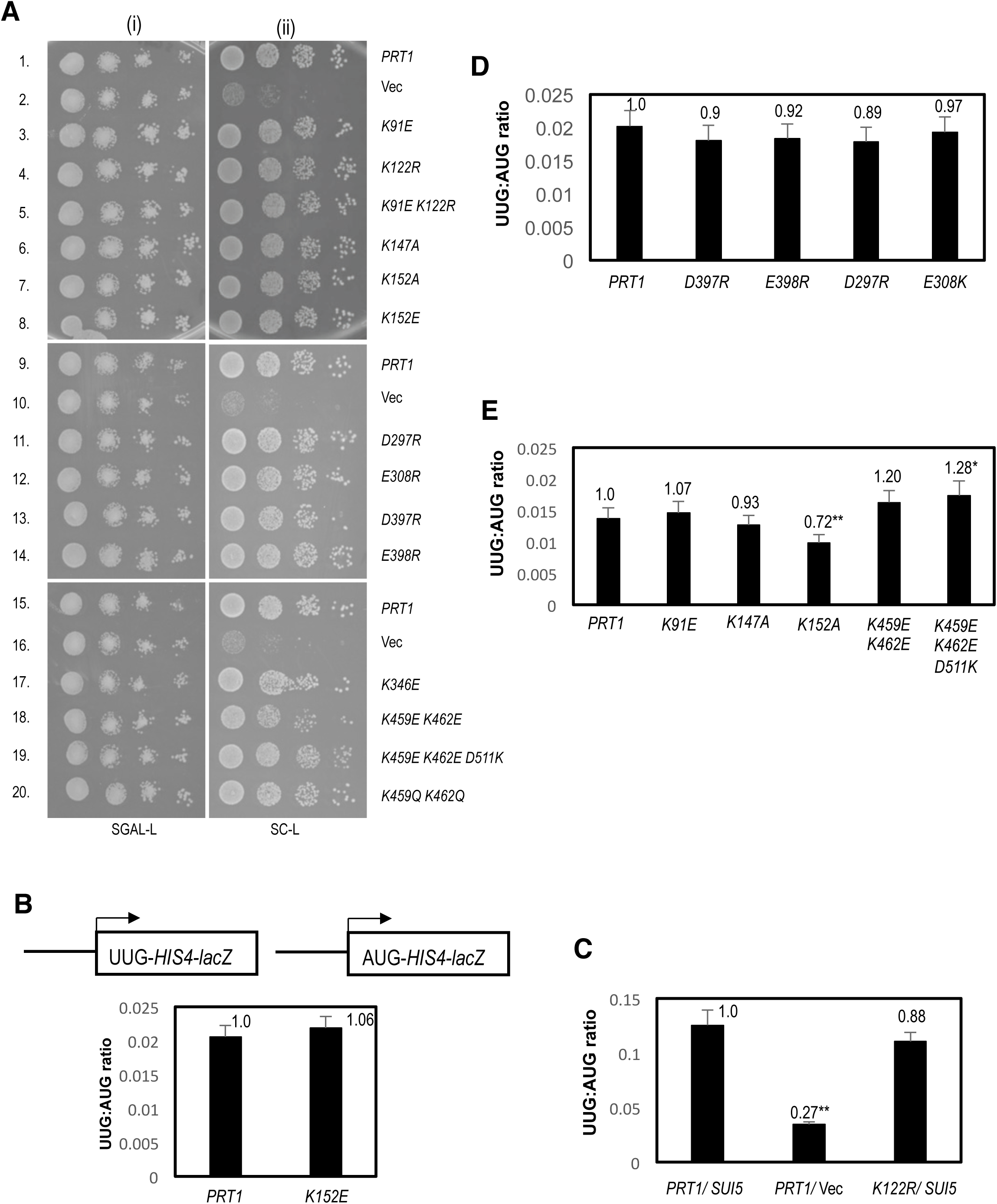
Supporting phenotypic analysis of *prt1* alleles. (A) Effects of *prt1* alleles on growth in complete medium in otherwise WT cells. Serial dilutions of the *P_GAL_-PRT1 his4-301* strain HD3607 harboring the indicated plasmid-borne *PRT1* alleles or empty vector (Vec) were spotted on SGAL-L or SC-L medium and incubated for 3-4 d at 30^0^ C. **(B, D, E) Effects of selected *prt1* alleles on the *HIS4-lacZ* UUG:AUG initiation ratio in otherwise WT cells.** Transformants of *P_GAL1_-PRT1 his4-301* strain HD3607 with the indicated *PRT1* alleles and *HIS4-lacZ* fusions with AUG or UUG start codons were cultured in SD+His+Trp, and analyzed exactly as in Figure 6B. **(C) *prt1-K112R* does not suppress the increased *HIS4-lacZ* UUG:AUG initiation ratio conferred by *SUI5*.** The *P_GAL_-PRT1 his4-301* strain HD3607 with the indicated plasmid-borne *PRT1* allele and harboring a single copy (sc) *SUI5* plasmid or empty vector (Vec), and *HIS4-lacZ* fusions with AUG or UUG start codons were analyzed exactly as in Figure 6B.

**Movie 1**. Movie showing the transition from py48S-open-eIF3 to py48S-closed-eIF3 PICs, in two different orientations and highlighting the subtle rearrangement of eIF3 elements at the subunit interface in this rearrangement.

**Table S1.** Summary of phenotypes of *PRT1* alleles.

| *PRT1* allele | Cell Growth at on SC | <sup>1</sup>Sui<sup>-</sup> | Suppression of *SUI5* His<sup>+</sup> | Suppression of *SUI5* Slg<sup>-</sup> at 37°C | *HIS4-lacZ* UUG:AUG in *SUI5* | Suppression of *SUI5* UUG:AUG |
| --- | --- | --- | --- | --- | --- | --- |
| WT | ++++ | none | None | none | 0.15±0.012 | none |
| *K91E* | ++++ | none | None | none | NA | NA |
| *K122R* | ++++ | Weak | None | none | 0.13±0.008 | none |
| *K91E*<br>*K122R* | ++++ | none | None | none | ND | ND |
| *K147A* | ++++ | none | Strong | Strong | 0.036±0.001 | Strong |
| *K152E* | ++++ | none | Strong | Strong | 0.033±0.005 | Strong |
| *K152A* | ++++ | none | Modest | strong | 0.082±0.009 | Modest |
| *D397R* | ++++ | none | Strong | Strong | 0.042±0.004 | Strong |
| *E398R* | ++++ | none | Strong | Strong | 0.050±0.006 | Strong |
| *D297R* | ++++ | none | None | Strong | 0.014±0.001 | Strong |
| *E308K* | ++++ | none | None | Strong | 0.006±0.002 | Strong |
| *K346E* | ++++ | Weak | none* | none* | ND | ND |
| *K459E*<br>*K462E* | ++ | none | Strong | Strong<sup>-</sup> | 0.039±0.005 | Strong |
| *K459Q*<br>*K462Q* | ++++ | ND | Strong | Strong | 0.075±0.010 | Modest |
| *K459E*<br>*K462E*<br>*D511K* | +++ | none | None | none | 0.15±0.009 | none |

<sup>1</sup>as judged by *HIS4-lacZ* UUG:AUG initiation ratio in strains lacking *SUI5*: none indicates an essentially WT ratio; weak indicates an ≈2-fold increase compared to the WT ratio.
NA: Not Applicable. Extreme Slg<sup>-</sup> in the presence of *SUI5* prevented introduction of *HIS4-lacZ* reporter plasmids.
ND: Not Determined
\*Data not shown in figures

**Table S2.**
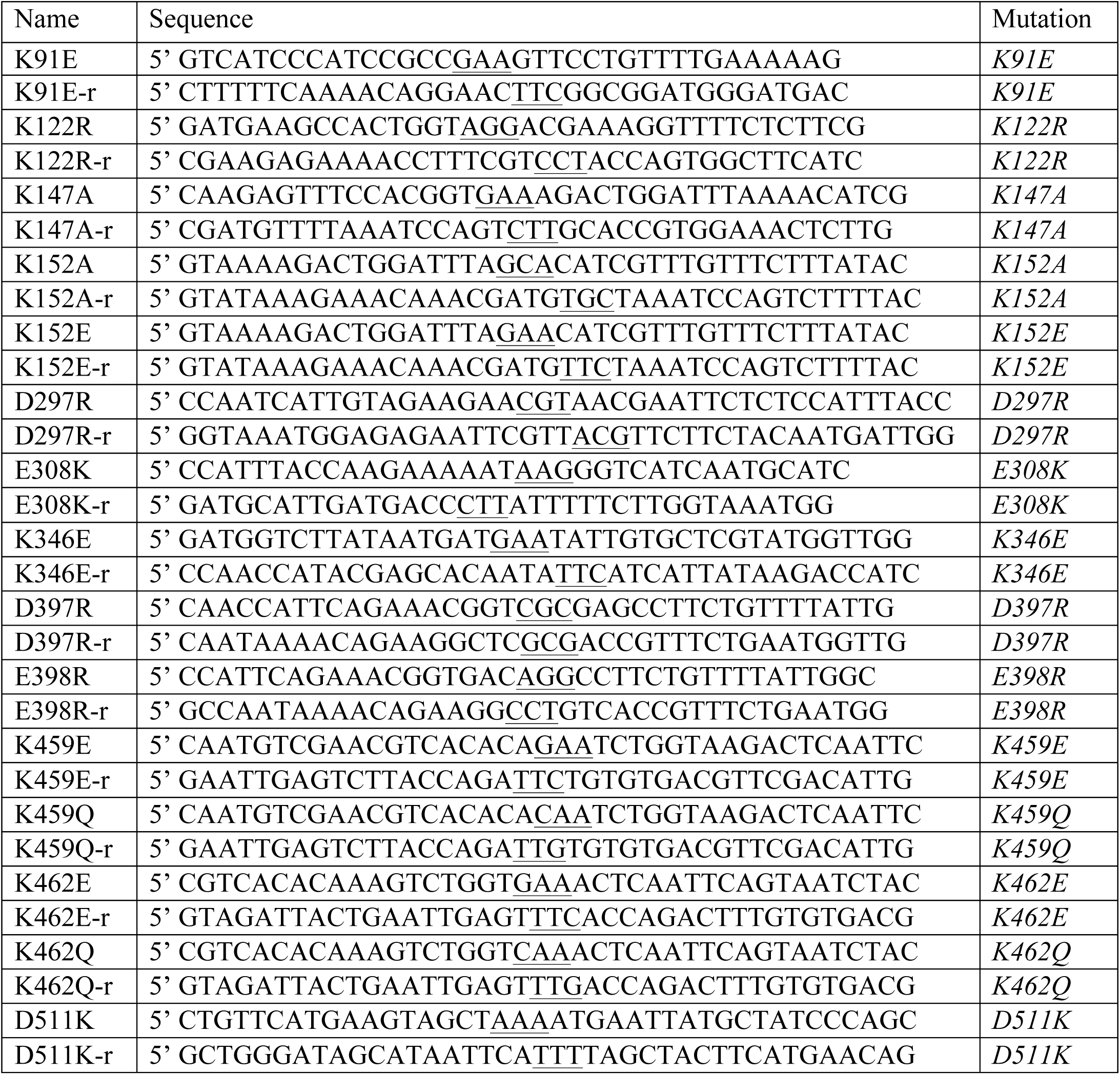
Primers used for mutagenesis (with mutated positions underlined)

**Table S3.**
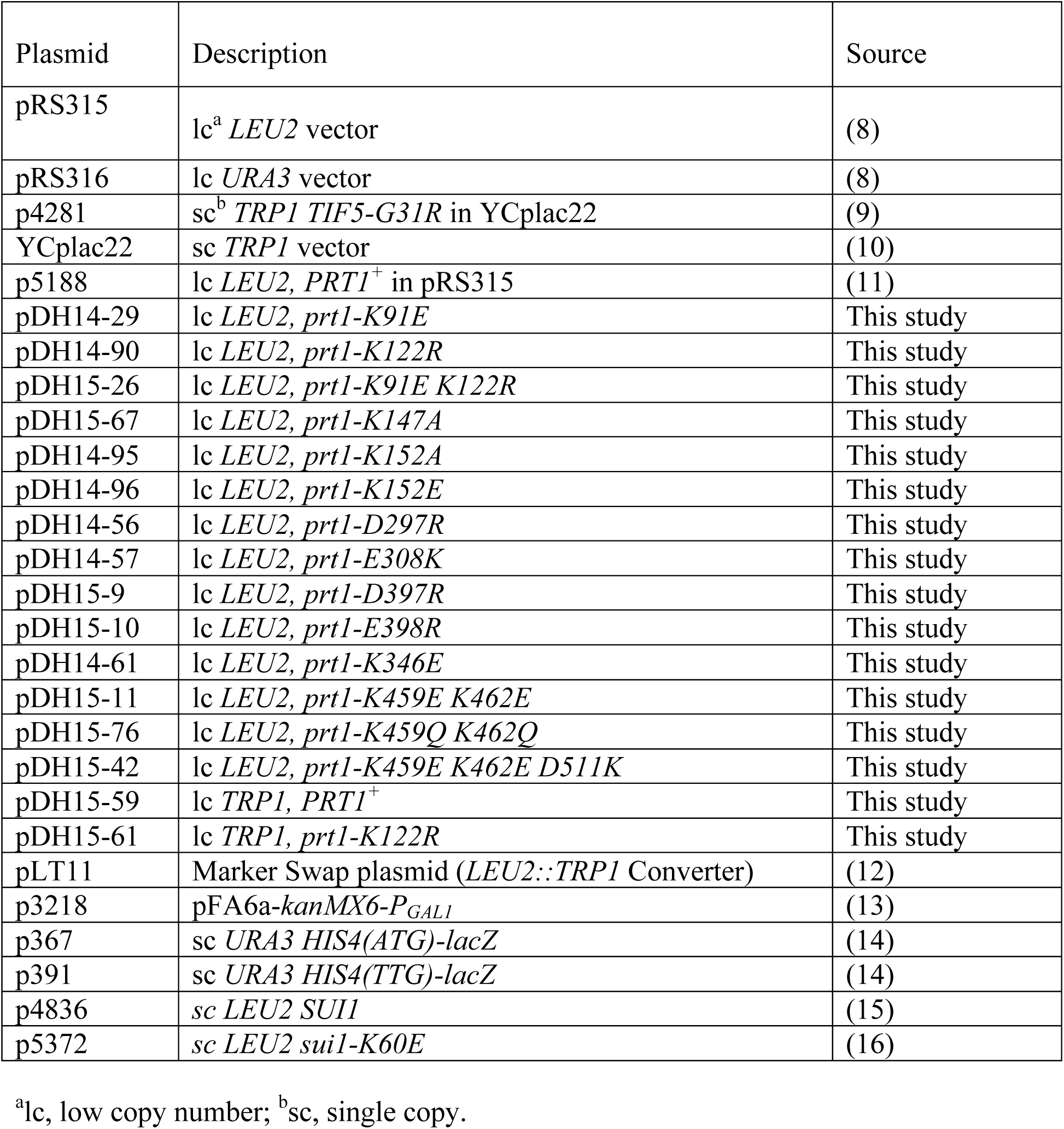
Plasmids employed in this work.

**Table S4.** Yeast strains employed in this work.

| Strain | Genotype | Source |
| --- | --- | --- |
| H2995 | *MATa ura3-52 trp1-63 leu2-3, 112 his4-301(ACG)* | (9) |
| HD3607 | *MATa ura3-52 trp1-63 leu2-3, 112 his4-301(ACG) P<sub>GAL</sub>-PRT1::KanMX* | This study |
| HD3648 | *MATa ura3-52 trp1-63 leu2-3, 112 his4-301(ACG) P<sub>GAL</sub>-PRT1::KanMX p5188 [lc LEU2 PRT1 in pRS315]* | This study |
| HD3687 | *MATa ura3-52 trp1-63 leu2-3, 112 his4-301(ACG) P<sub>GAL</sub>-PRT1::KanMX pRS315 [lc LEU2]* | This study |
| HD3649 | *MATa ura3-52 trp1-63 leu2-3, 112 his4-301(ACG) P<sub>GAL</sub>-PRT1::KanMX pDH14-29 [lc LEU2 prt1-K91E in pRS315]* | This study |
| HD3848 | *MATa ura3-52 trp1-63 leu2-3, 112 his4-301(ACG) P<sub>GAL</sub>-PRT1::KanMX pDH14-90 [lc LEU2 prt1-K122R in pRS315]* | This study |
| HD4059 | *MATa ura3-52 trp1-63 leu2-3, 112 his4-301(ACG) P<sub>GAL</sub>-PRT1::KanMX pDH15-26 [lc LEU2 prt1-K91E K122R in pRS315]* | This study |
| HD4262 | *MATa ura3-52 trp1-63 leu2-3, 112 his4-301(ACG) P<sub>GAL</sub>-PRT1::KanMX pDH15-67 [lc LEU2 prt1-K147A in pRS315]* | This study |
| HD4256 | *MATa ura3-52 trp1-63 leu2-3, 112 his4-301(ACG) P<sub>GAL</sub>-PRT1::KanMX pDH14-95 [lc LEU2 prt1-K152A in pRS315]* | This study |
| HD3850 | *MATa ura3-52 trp1-63 leu2-3, 112 his4-301(ACG) P<sub>GAL</sub>-PRT1::KanMX pDH14-96 [lc LEU2 prt1-K152E in pRS315]* | This study |
| HD3668 | *MATa ura3-52 trp1-63 leu2-3, 112 his4-301(ACG) P<sub>GAL</sub>-PRT1::KanMX pDH14-56 [lc LEU2 prt1-D297R in pRS315]* | This study |
| HD3669 | *MATa ura3-52 trp1-63 leu2-3, 112 his4-301(ACG) P<sub>GAL</sub>-PRT1::KanMX pDH14-57 [lc LEU2 prt1-E308K in pRS315]* | This study |
| HD3861 | *MATa ura3-52 trp1-63 leu2-3, 112 his4-301(ACG) P<sub>GAL</sub>-PRT1::KanMX pDH15-9 [lc LEU2 prt1-D397R in pRS315]* | This study |
| HD3862 | *MATa ura3-52 trp1-63 leu2-3, 112 his4-301(ACG) P<sub>GAL</sub>-PRT1::KanMX pDH15-10 [lc LEU2 prt1-E398R in pRS315]* | This study |
| HD3672 | *MATa ura3-52 trp1-63 leu2-3, 112 his4-301(ACG) P<sub>GAL</sub>-PRT1::KanMX pDH14-61 [lc LEU2 prt1-K346E in pRS315]* | This study |
| HD3955 | *MATa ura3-52 trp1-63 leu2-3, 112 his4-301(ACG) P<sub>GAL</sub>-PRT1::KanMX pDH15-11 [lc LEU2 prt1-K459E K462E in pRS315]* | This study |
| HD4263 | *MATa ura3-52 trp1-63 leu2-3, 112 his4-301(ACG)*<br>*P<sub>GAL</sub>-PRT1::KanMX* pDH15-76 [lc *LEU2 prt1-K459Q*<br>*K462Q* in pRS315] | This study |
| HD4075 | *MATa ura3-52 trp1-63 leu2-3, 112 his4-301(ACG)*<br>*P<sub>GAL</sub>-PRT1::KanMX* pDH15-42 [lc *LEU2 prt1-K459E*<br>*K462E D511K* in pRS315] | This study |
| HD4081 | *MATa ura3-52 trp1-63 leu2-3, 112 his4-301(ACG)*<br>*P<sub>GAL</sub>-PRT1::KanMX* p5188 [lc *LEU2 PRT1<sup>+</sup>* in pRS315]<br>p4281 [sc *TRP1 SUI5*] | This study |
| HD4082 | *MATa ura3-52 trp1-63 leu2-3, 112 his4-301(ACG)*<br>*P<sub>GAL</sub>-PRT1::KanMX* p5188 [lc *LEU2 PRT1<sup>+</sup>* in pRS315]<br>Ycplac22 [sc *TRP1*] | This study |
| HD4269 | *MATa ura3-52 trp1-63 leu2-3, 112 his4-301(ACG)*<br>*P<sub>GAL</sub>-PRT1::KanMX* pDH15-67 [lc *LEU2 prt1-K147A* in<br>pRS315] p4281 [sc *TRP1 SUI5*] | This study |
| HD3995 | *MATa ura3-52 trp1-63 leu2-3, 112 his4-301(ACG)*<br>*P<sub>GAL</sub>-PRT1::KanMX* pDH14-96 [lc *LEU2 prt1-K152E* in<br>pRS315] p4281 [sc *TRP1 SUI5*] | This study |
| HD4257 | *MATa ura3-52 trp1-63 leu2-3, 112 his4-301(ACG)*<br>*P<sub>GAL</sub>-PRT1::KanMX* pDH14-95 [lc *LEU2 prt1-K152A* in<br>pRS315] p4281 [sc *TRP1 SUI5*] | This study |
| HD4280 | *MATa ura3-52 trp1-63 leu2-3, 112 his4-301(ACG)*<br>*P<sub>GAL</sub>-PRT1::KanMX* pDH15-9 [lc *LEU2 prt1-D397R* in<br>pRS315] p4281 [sc *TRP1 SUI5*] | This study |
| HD4281 | *MATa ura3-52 trp1-63 leu2-3, 112 his4-301(ACG)*<br>*P<sub>GAL</sub>-PRT1::KanMX* pDH15-10 [lc *LEU2 prt1-E398R* in<br>pRS315] p4281 [sc *TRP1 SUI5*] | This study |
| HD3749 | *MATa ura3-52 trp1-63 leu2-3, 112 his4-301(ACG)*<br>*P<sub>GAL</sub>-PRT1::KanMX* pDH14-56 [lc *LEU2 prt1-D297R* in<br>pRS315] p4281 [sc *TRP1 SUI5*] | This study |
| HD4279 | *MATa ura3-52 trp1-63 leu2-3, 112 his4-301(ACG)*<br>*P<sub>GAL</sub>-PRT1::KanMX* pDH14-57 [lc *LEU2 prt1-E308K* in<br>pRS315] p4281 [sc *TRP1 SUI5*] | This study |
| HD4012 | *MATa ura3-52 trp1-63 leu2-3, 112 his4-301(ACG)*<br>*P<sub>GAL</sub>-PRT1::KanMX* pDH15-11 [lc *LEU2 prt1-K459E*<br>*K462E* in pRS315] p4281 [sc *TRP1 SUI5*] | This study |
| HD4268 | *MATa ura3-52 trp1-63 leu2-3, 112 his4-301(ACG)*<br>*P<sub>GAL</sub>-PRT1::KanMX* pDH15-76 [lc *LEU2 prt1-K459Q*<br>*K462Q* in pRS315] p4281 [sc *TRP1 SUI5*] | This study |
| HD4009 | *MATa ura3-52 trp1-63 leu2-3, 112 his4-301(ACG)*<br>*P<sub>GAL</sub>-PRT1::KanMX* pDH15-42 [lc *LEU2 prt1-K459E*<br>*K462E D511K* in pRS315] p4281 [sc *TRP1 SUI5*] | This study |
| HD3993 | *MATa ura3-52 trp1-63 leu2-3, 112 his4-301(ACG)* *P<sub>GAL</sub>-*<br>*PRT1::KanMX* pDH14-90 [lc *LEU2 prt1-K122R* in<br>pRS315] p4281 [sc *TRP1 SUI5*] | This study |
| H3956 | *MATa ura3-52 leu2-3,112 trp1-Δ63 his4-303(AUU) sui1Δ::hisG p1200 [SUI1, URA3 CEN4]* | This study |
| HD4053 | *MATa ura3-52 leu2-3,112 trp1-Δ63 his4-303(AUU) P<sub>GAL</sub>-PRT1::KanMX sui1Δ::hisG p1200 [sc SUI1, URA3 CEN4].* | This study |
| HD4108 | *MATa ura3-52 leu2-3,112 trp1-Δ63 his4-303(AUU) P<sub>GAL</sub>-PRT1 sui1Δ::hisG p4836 [sc SUI1, LEU2 CEN4].* | This study |
| HD4109 | *(MATa ura3-52 leu2-3,112 trp1-Δ63 his4-303(AUU) P<sub>GAL</sub>-PRT1::KanMX sui1Δ::hisG p5372 [sc sui1-K60E, LEU2 CEN4].* | This study |
| HD4192 | *MATa ura3-52 leu2-3,112 trp1-Δ63 his4-303(AUU) P<sub>GAL</sub>-PRT1::KanMX sui1Δ::hisG p4836 [SUI1, LEU2 CEN4] pDH15-59 [lc PRT1 TRP1].* | This study |
| HD4193 | *MATa ura3-52 leu2-3,112 trp1-Δ63 his4-303(AUU) P<sub>GAL</sub>-PRT1::KanMX sui1Δ::hisG p5372 [sui1-K60E, LEU2 CEN4] pDH15-59 [lc PRT1 TRP1]* | This study |
| HD4196 | *MATa ura3-52 leu2-3,112 trp1-Δ63 his4-303(AUU) P<sub>GAL</sub>-PRT1::KanMX sui1Δ::hisG p4836 [SUI1, LEU2 CEN4] pDH15-61 [lc prt1-K122R TRP1]* | This study |
| HD4197 | *MATa ura3-52 leu2-3,112 trp1-Δ63 his4-303(AUU) P<sub>GAL</sub>-PRT1::KanMX sui1Δ::hisG p5372 [sui1-K60E, LEU2 CEN4] pDH15-61[lc prt1-K122R TRP1]* | This study |

## References

Acker, M. G., Kolitz, S. E., Mitchell, S. F., Nanda, J. S., and Lorsch, J. R. (2007). Reconstitution of yeast translation initiation. Methods Enzymol 430, 111–145.

Amunts, A., Brown, A., Bai, X. C., Llácer, J. L., Hussain, T., Emsley, P., Long, F., Murshudov, G., Scheres, S. H. W., and Ramakrishnan, V. (2014). Structure of the yeast mitochondrial large ribosomal subunit. Science 343, 1485–1489.

Aylett, C. H., and Ban, N. (2017). Eukaryotic aspects of translation initiation brought into focus. Philos Trans R Soc Lond B Biol Sci 372,

Aylett, C. H., Boehringer, D., Erzberger, J. P., Schaefer, T., and Ban, N. (2015). Structure of a yeast 40S-eIF1-eIF1A-eIF3-eIF3j initiation complex. Nat Struct Mol Biol 22, 269–271.

Bai, X. C., Fernandez, I. S., McMullan, G., and Scheres, S. H. (2013). Ribosome structures to near-atomic resolution from thirty thousand cryo-EM particles. Elife 2, e00461.

Bai, X. C., Rajendra, E., Yang, G., Shi, Y., and Scheres, S. H. (2015). Sampling the conformational space of the catalytic subunit of human γ-secretase. Elife 4,

Brown, A., Long, F., Nicholls, R. A., Toots, J., Emsley, P., and Murshudov, G. (2015). Tools for macromolecular model building and refinement into electron cryo-microscopy reconstructions. Acta Crystallogr D Biol Crystallogr 71, 136–153.

Chen, V. B., Arendall, W. B., Headd, J. J., Keedy, D. A., Immormino, R. M., Kapral, G. J., Murray, L. W., Richardson, J. S., and Richardson, D. C. (2010). MolProbity: all-atom structure validation for macromolecular crystallography. Acta Crystallogr D Biol Crystallogr 66, 12–21.

Chiu, W. L., Wagner, S., Herrmannová, A., Burela, L., Zhang, F., Saini, A. K., Valásek, L., and Hinnebusch, A. G. (2010). The C-terminal region of eukaryotic translation initiation factor 3a (eIF3a) promotes mRNA recruitment, scanning, and, together with eIF3j and the eIF3b RNA recognition motif, selection of AUG start codons. Mol Cell Biol 30, 4415–4434.

Cross, F. R. (1997). Marker swap’ plasmids: convenient tools for budding yeast molecular genetics. Yeast 13, 647–653.

DeLano, W. L. (2006). The PyMOL Molecular Graphics System. http://www.pymol.org

des Georges, A., Dhote, V., Kuhn, L., Hellen, C. U., Pestova, T. V., Frank, J., and Hashem, Y. (2015). Structure of mammalian eIF3 in the context of the 43S preinitiation complex. Nature 525, 491–495.

Eliseev, B., Yeramala, L., Leitner, A., Karuppasamy, M., Raimondeau, E., Huard, K., Alkalaeva, E., Aebersold, R., and Schaffitzel, C. (2018). Structure of a human cap-dependent 48S translation pre-initiation complex. Nucleic Acids Res 46, 2678–2689.

Emsley, P., Lohkamp, B., Scott, W. G., and Cowtan, K. (2010). Features and development of Coot. Acta Crystallogr D Biol Crystallogr 66, 486–501.

Erzberger, J. P., Stengel, F., Pellarin, R., Zhang, S., Schaefer, T., Aylett, C. H. S., Cimermančič, P., Boehringer, D., Sali, A., Aebersold, R., and Ban, N. (2014). Molecular architecture of the 40S⋅eIF1⋅eIF3 translation initiation complex. Cell 158, 1123–1135.

Fernández, I. S., Bai, X. C., Murshudov, G., Scheres, S. H., and Ramakrishnan, V. (2014). Initiation of translation by cricket paralysis virus IRES requires its translocation in the ribosome. Cell 157, 823–831.

Hashem, Y., des Georges, A., Dhote, V., Langlois, R., Liao, H. Y., Grassucci, R. A., Hellen, C. U., Pestova, T. V., and Frank, J. (2013). Structure of the mammalian ribosomal 43S preinitiation complex bound to the scanning factor DHX29. Cell 153, 1108–1119.

Herrmannová, A., Daujotyte, D., Yang, J. C., Cuchalová, L., Gorrec, F., Wagner, S., Dányi, I., Lukavsky, P. J., and Valásek, L. S. (2012). Structural analysis of an eIF3 subcomplex reveals conserved interactions required for a stable and proper translation pre-initiation complex assembly. Nucleic Acids Res 40, 2294–2311.

Heuer, A., Gerovac, M., Schmidt, C., Trowitzsch, S., Preis, A., Kötter, P., Berninghausen, O., Becker, T., Beckmann, R., and Tampé, R. (2017). Structure of the 40S-ABCE1 post-splitting complex in ribosome recycling and translation initiation. Nat Struct Mol Biol 24, 453–460.

Hinnebusch, A. G. (2014). The scanning mechanism of eukaryotic translation initiation. Annu Rev Biochem 83, 779–812.

Hinnebusch, A. G. (2017). Structural Insights into the Mechanism of Scanning and Start Codon Recognition in Eukaryotic Translation Initiation. Trends Biochem Sci 42, 589–611.

Huang, H. K., Yoon, H., Hannig, E. M., and Donahue, T. F. (1997). GTP hydrolysis controls stringent selection of the AUG start codon during translation initiation in Saccharomyces cerevisiae. Genes Dev 11, 2396–2413.

Hussain, T., Llácer, J. L., Fernández, I. S., Munoz, A., Martin-Marcos, P., Savva, C. G., Lorsch, J. R., Hinnebusch, A. G., and Ramakrishnan, V. (2014). Structural changes enable start codon recognition by the eukaryotic translation initiation complex. Cell 159, 597–607.

Hussain, T., Llácer, J. L., Wimberly, B. T., Kieft, J. S., and Ramakrishnan, V. (2016). Large-Scale Movements of IF3 and tRNA during Bacterial Translation Initiation. Cell 167, 133–144.e13.

Kashiwagi, K., Yokoyama, T., Nishimoto, M., Takahashi, M., Sakamoto, A., Yonemochi, M., Shirouzu, M., and Ito, T. (2019). Structural basis for eIF2B inhibition in integrated stress response. Science 364, 495–499.

Kenner, L. R., Anand, A. A., Nguyen, H. C., Myasnikov, A. G., Klose, C. J., McGeever, L. A., Tsai, J. C., Miller-Vedam, L. E., Walter, P., and Frost, A. (2019). eIF2B-catalyzed nucleotide exchange and phosphoregulation by the integrated stress response. Science 364, 491–495.

Khoshnevis, S., Gunišová, S., Vlčková, V., Kouba, T., Neumann, P., Beznosková, P., Ficner, R., and Valášek, L. S. (2014). Structural integrity of the PCI domain of eIF3a/TIF32 is required for mRNA recruitment to the 43S pre-initiation complexes. Nucleic Acids Res 42, 4123–4139.

Krissinel, E., and Henrick, K. (2007). Inference of macromolecular assemblies from crystalline state. J Mol Biol 372, 774–797.

Kucukelbir, A., Sigworth, F. J., and Tagare, H. D. (2014). Quantifying the local resolution of cryo-EM density maps. Nat Methods 11, 63–65.

Li, X., Mooney, P., Zheng, S., Booth, C. R., Braunfeld, M. B., Gubbens, S., Agard, D. A., and Cheng, Y. (2013). Electron counting and beam-induced motion correction enable near-atomic-resolution single-particle cryo-EM. Nat Methods 10, 584–590.

Llácer, J. L., Hussain, T., Marler, L., Aitken, C. E., Thakur, A., Lorsch, J. R., Hinnebusch, A. G., and Ramakrishnan, V. (2015). Conformational Differences between Open and Closed States of the Eukaryotic Translation Initiation Complex. Mol Cell 59, 399–412.

Llácer, J. L., Hussain, T., Saini, A. K., Nanda, J. S., Kaur, S., Gordiyenko, Y., Kumar, R., Hinnebusch, A. G., Lorsch, J. R., and Ramakrishnan, V. (2018). Translational initiation factor eIF5 replaces eIF1 on the 40S ribosomal subunit to promote start-codon recognition. Elife 7,

Mancera-Martínez, E., Brito Querido, J., Valasek, L. S., Simonetti, A., and Hashem, Y. (2017). ABCE1: A special factor that orchestrates translation at the crossroad between recycling and initiation. RNA Biol 14, 1279–1285.

Martin-Marcos, P., Cheung, Y. N., and Hinnebusch, A. G. (2011). Functional elements in initiation factors 1, 1A, and 2β discriminate against poor AUG context and non-AUG start codons. Mol Cell Biol 31, 4814–4831.

Martin-Marcos, P., Nanda, J., Luna, R. E., Wagner, G., Lorsch, J. R., and Hinnebusch, A. G. (2013). β-Hairpin loop of eukaryotic initiation factor 1 (eIF1) mediates 40 S ribosome binding to regulate initiator tRNA(Met) recruitment and accuracy of AUG selection in vivo. J Biol Chem 288, 27546–27562.

Martin-Marcos, P., Nanda, J. S., Luna, R. E., Zhang, F., Saini, A. K., Cherkasova, V. A., Wagner, G., Lorsch, J. R., and Hinnebusch, A. G. (2014). Enhanced eIF1 binding to the 40S ribosome impedes conformational rearrangements of the preinitiation complex and elevates initiation accuracy. RNA 20, 150–167.

Mishra, R. K., Datey, A., and Hussain, T. (2019). mRNA Recruiting eIF4 Factors Involved in Protein Synthesis and Its Regulation. Biochemistry

Mitchell, S. F., Walker, S. E., Algire, M. A., Park, E. H., Hinnebusch, A. G., and Lorsch, J. R. (2010). The 5’-7-methylguanosine cap on eukaryotic mRNAs serves both to stimulate canonical translation initiation and to block an alternative pathway. Mol Cell 39, 950–962.

Moehle, C. M., and Hinnebusch, A. G. (1991). Association of RAP1 binding sites with stringent control of ribosomal protein gene transcription in Saccharomyces cerevisiae. Mol Cell Biol 11, 2723–2735.

Pelletier, J., and Sonenberg, N. (2019). The Organizing Principles of Eukaryotic Ribosome Recruitment. Annu Rev Biochem 88, 307–335.

Pettersen, E. F., Goddard, T. D., Huang, C. C., Couch, G. S., Greenblatt, D. M., Meng, E. C., and Ferrin, T. E. (2004). UCSF Chimera--a visualization system for exploratory research and analysis. J Comput Chem 25, 1605–1612.

Ramlaul, K., Palmer, C. M., and Aylett, C. H. S. (2019). A Local Agreement Filtering Algorithm for Transmission EM Reconstructions. J Struct Biol 205, 30–40.

Reibarkh, M., Yamamoto, Y., Singh, C. R., del Rio, F., Fahmy, A., Lee, B., Luna, R. E., Ii, M., Wagner, G., and Asano, K. (2008). Eukaryotic initiation factor (eIF) 1 carries two distinct eIF5-binding faces important for multifactor assembly and AUG selection. J Biol Chem 283, 1094–1103.

Rosenthal, P. B., and Henderson, R. (2003). Optimal determination of particle orientation, absolute hand, and contrast loss in single-particle electron cryomicroscopy. J Mol Biol 333, 721–745.

Saini, A. K., Nanda, J. S., Martin-Marcos, P., Dong, J., Zhang, F., Bhardwaj, M., Lorsch, J. R., and Hinnebusch, A. G. (2014). Eukaryotic translation initiation factor eIF5 promotes the accuracy of start codon recognition by regulating Pi release and conformational transitions of the preinitiation complex. Nucleic Acids Res 42, 9623–9640.

Scheres, S. H. (2012). RELION: implementation of a Bayesian approach to cryo-EM structure determination. J Struct Biol 180, 519–530.

Scheres, S. H. (2015). Semi-automated selection of cryo-EM particles in RELION-1.3. J Struct Biol 189, 114–122.

Scheres, S. H., and Chen, S. (2012). Prevention of overfitting in cryo-EM structure determination. Nat Methods 9, 853–854.

Simonetti, A., Brito Querido, J., Myasnikov, A. G., Mancera-Martinez, E., Renaud, A., Kuhn, L., and Hashem, Y. (2016). eIF3 Peripheral Subunits Rearrangement after mRNA Binding and Start-Codon Recognition. Mol Cell 63, 206–217.

Szamecz, B., Rutkai, E., Cuchalová, L., Munzarová, V., Herrmannová, A., Nielsen, K. H., Burela, L., Hinnebusch, A. G., and Valásek, L. (2008). eIF3a cooperates with sequences 5’ of uORF1 to promote resumption of scanning by post-termination ribosomes for reinitiation on GCN4 mRNA. Genes Dev 22, 2414–2425.

Tang, G., Peng, L., Baldwin, P. R., Mann, D. S., Jiang, W., Rees, I., and Ludtke, S. J. (2007). EMAN2: an extensible image processing suite for electron microscopy. J Struct Biol 157, 38–46.

Thakur, A., and Hinnebusch, A. G. (2018). eIF1 Loop 2 interactions with Met-tRNA. Proc Natl Acad Sci U S A 115, E4159–E4168.

Valášek, L. S., Zeman, J., Wagner, S., Beznosková, P., Pavlíková, Z., Mohammad, M. P., Hronová, V., Herrmannová, A., Hashem, Y., and Gunišová, S. (2017). Embraced by eIF3: structural and functional insights into the roles of eIF3 across the translation cycle. Nucleic Acids Res 45, 10948–10968.

Zeman, J., Itoh, Y., Kukačka, Z., Rosůlek, M., Kavan, D., Kouba, T., Jansen, M. E., Mohammad, M. P., Novák, P., and Valášek, L. S. (2019). Binding of eIF3 in complex with eIF5 and eIF1 to the 40S ribosomal subunit is accompanied by dramatic structural changes. Nucleic Acids Res 47, 8282–8300.

Zhang, K. (2016). Gctf: Real-time CTF determination and correction. J Struct Biol 193, 1–12.

